# The interplay between glucose and aromatic compound regulation by two IclR-type transcription factors, LigR1 and LigR2, in *Pseudomonas putida* KT2440

**DOI:** 10.1101/2025.08.13.670189

**Authors:** Elina Kadriu, Sophie Qin, Stephanie Prezioso, Dinesh Christendat

## Abstract

The rhizosphere is a hotspot of microbial activity where plants release a diverse array of aromatic compounds, including shikimate pathway intermediates and monolignols. *Pseudomonas putida* KT2440, renowned for its metabolic versatility in this niche, uses largely uncharacterized regulatory and enzymatic strategies to utilize these compounds. We investigated two IclR-type transcriptional regulators, LigR1 and LigR2, that control expression of the uncharacterized *lig1* and *lig2* operons. We demonstrate that *ligR1* deletion caused growth defects on glucose and 4-hydroxybenzoate accompanied by cell elongation and aggregation. Structural and functional analyses reveal that LigR1 and LigR2 activate the *lig1* operon but repress the *lig2* operon. LigR1 binding of 4-hydroxybenzoate induced repression by triggering tetramerization and increasing DNA-binding activity. In contrast, LigR2 responded to quinate, protocatechuate and 4-hydroxybenzoate to potently induce *lig2* operon expression by relieving repression. While both operons cooperate in metabolizing these compounds, we propose the *lig1* operon mediates influx through its major facilitator superfamily (MFS) transporter (PP_2604), whereas the *lig2* operon catalyzes breakdown via a protocatechuate intermediate and the meta-cleavage pathway, supplying oxaloacetate to the TCA cycle. Importantly, we show that *P. putida* repurposes shikimate pathway intermediates for energy production. These findings challenge the canonical biosynthetic view of the shikimate pathway and redefine the metabolic flexibility of soil pseudomonads. We reveal a novel mechanism enabling *P. putida* to thrive in the chemically complex rhizosphere and open new avenues for exploring alternate roles of the shikimate pathway, emphasizing transcriptional regulators as tools to deconvolute complex metabolic landscapes.

**Highlights:**

- LigR1 and LigR2 transcriptionally regulate the *lig1* and *lig2* operons
- *Lig1* operon is required for import of glucose and shikimate-derived compounds
- *Lig2* operon metabolizes shikimate pathway compounds
- Dysregulated LigR1/LigR2 expression impacts bacterial physiology

**Graphical Abstract:** 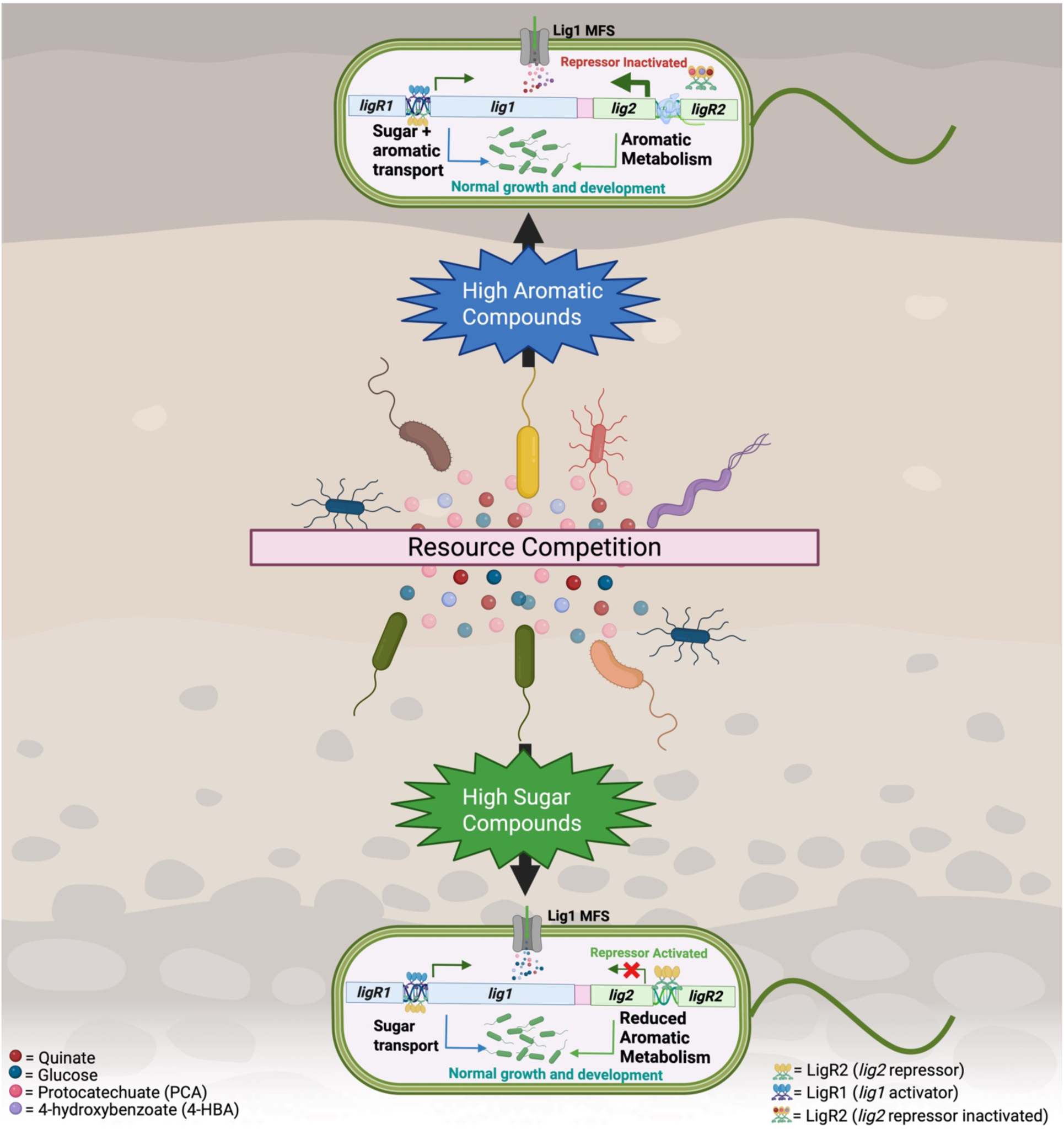

## Introduction

The rhizosphere is a dynamic and metabolically complex region surrounding plant roots where microorganisms compete for both nutrients and space^1^. The nutritional landscape within this niche fluctuates continuously, is influenced by changes in carbon source composition and availability, and shaped by plant species, and soil type^2,3^. *Pseudomonas putida* KT2440 thrives amongst the microbes inhabiting the rhizosphere, due to its highly versatile metabolic networks^4^. Studies have shown that *P. putida* KT2440 metabolizes carbon compounds in a hierarchical order, preferring organic acids first, followed by amino acids, hexose sugars including glucose, and lastly aromatic compounds^5^. In contrast, other species such as *P. putida* M2 and *P. putida* CSV86 preferentially utilize aromatic compounds even with an overabundance of sugars^5,6^.

The metabolism of sugars and aromatics by *P. putida* is tightly regulated via a suite of transcriptional regulators, which act locally to control gene expression^7–9^. Additionally, *P. putida* KT2440 employs a well-characterized global regulatory mechanism: the carbon catabolite repression (CCR) system controlled mainly by the Crc and Hfq regulators^10,11^. The CCR system represses translation of proteins encoded by genes involved in the metabolism of non-preferred carbon sources until preferred substrates are available, thereby optimizing resource allocation^10,12^. The ability to rapidly reprogram its transcriptional landscape gives *P. putida* KT2440 a competitive edge in colonizing and surviving in environments with fluctuating carbon availability^4^.

In *P. putida* KT2440, glucose and aromatic compounds are metabolized through distinct biochemical pathways that ultimately converge at the TCA cycle for energy production^9,13^. Glucose utilization occurs predominantly via the Entner-Doudoroff (ED) pathway, although the Embden-Meyerhof-Parnas (EMP) and pentose phosphate (PP) pathways are also present^13^. Glucose first enters the periplasmic space through the OprB porin in the outer membrane, after which it may be directly transported into the cytoplasm by the GtsABCD ABC transporter or oxidized into gluconate and 2-ketogluconate via the enzymes glucose dehydrogenase (Gcd) and gluconate oxidase (Gad), respectively^13–15^. Once inside the cytoplasm, glucose, gluconate, and 2-ketogluconate are converted into 6-phosphogluconate (6PG) through distinct enzymatic routes^13,15^. Glucose follows a pathway involving glucokinase (Glk), glucose 6-phosphate dehydrogenase (Zwf), and 6-phosphogluconolactonase (Pgl), while gluconate is phosphorylated by gluconate kinase (GnuK), and 2-ketogluconate is processed by 2-ketogluconate kinase (KguK) and then reduced by 2-ketogluconate-6P reductase (KguD)^13,15^. The resulting 6PG is dehydrated by 6-phosphogluconate dehydratase (Edd) to form 2-keto-3-deoxy-6-phosphogluconate (KDPG), which is then cleaved by 2-keto-3-deoxy-6P-gluconate aldolase (Eda) into pyruvate and glyceraldehyde-3-phosphate (G3P). G3P can enter into the EMP pathway to form phosphoenolpyruvate (PEP) and pyruvate. The pyruvate from the ED and EMP pathways are converted into oxaloacetate, which enters the TCA cycle^13,15^.The remaining PEP byproduct from the EMP pathway combines with erythrose-4-phosphate from the PP pathway to form the initial intermediate of the shikimate pathway. This finally leads to chorismate biosynthesis and the downstream production of the three aromatic amino acids: tryptophan, tyrosine, and phenylalanine^16,17^.

The genes involved in glucose metabolism are organized into distinct operons that are subjected to tightly coordinated regulation^8^. The *zwf*, *pgl*, and *eda* genes are transcribed divergently from the *hexR* transcriptional regulator^8^. Meanwhile, *edd*, *glk*, and *gts* genes, along with their regulators *gltR2* and *gltS*, are transcribed divergently from *gap-1*, which encodes the EMP pathway enzyme glyceraldehyde-3-phosphate dehydrogenase^8^. The *gnuK* and *gltP* genes (for gluconate utilization) are oriented with the *gntR* regulator, and the *kguK*, *kguD*, and *kguT* genes (for 2-ketogluconate metabolism) are co-oriented with the *ptxS* regulator^8,18^. These transcription factors—HexR, GltR/GltS, GntR, and PtxS—function as either repressors (HexR, GntR, PtxS) or activators (GltR/GltS) and contain both an N-terminal DNA-binding domain and a C-terminal effector-binding domain that responds to intermediates of the ED pathway^8^.

For aromatic compound metabolism, *P. putida* KT2440 employs three central pathways: β-ketoadipate, homogentisate, and phenylacetate^9^. The β-ketoadipate pathway is divided into two branches: the protocatechuate branch, which processes lignin-derived compounds (e.g., coniferyl alcohol, ferulate, caffeate, cinnamate), and the catechol branch, which degrades benzoate-like compounds^9^. Both branches converge at β-ketoadipate, which is further cleaved into acetyl-CoA and succinyl-CoA^9^. The homogentisate pathway catabolizes phenylalanine and tyrosine into homogentisate, which is then metabolized into fumarate and acetoacetate^9^. These two pathways are transcriptionally controlled by IclR-family regulators: PcaR, which activates the *pca* operon in response to β-ketoadipate, and HmgR, which represses the *hmg* operon until relieved by homogentisate^19–22^.

IclR-type regulators form a diverse protein family controlling processes across bacteria and archaea, from carbon metabolism to pathogenesis^23^. Named after the glyoxylate bypass repressor in *E. coli*, IclRs regulate systems involved in biofilm formation, antibiotic biosynthesis, multidrug resistance, sugar metabolism, and aromatic compound degradation^23–29^. They are especially prominent in aromatic metabolism, where they regulate pathways for the breakdown of catechol, homogentisate, 4-hydroxybenzoate, and other plant-derived compounds^19,20,29–31^. This versatility has enabled their use in metabolic engineering and biosensor design^32–36^. Structurally, IclRs share a conserved architecture featuring an N-terminal helix-turn-helix DNA-binding domain and a C-terminal effector-binding domain, linked by a short α-helix^37^. Functioning as tetramers, these regulators undergo conformational shifts upon effector binding, distorting the DNA promoter into a characteristic ‘W’ shape^27,38^. Their regulatory effect such as activation, repression, or dual control depends on both the effector type and the operon function. For example, *E. coli* IclR activates the *aceBAK* operon with glyoxylate but represses it in the presence of pyruvate^39^. Many IclRs, such as the *Rhodococcus jostii* RHA1 TsdR, also autoregulate their own expression^40^. Mechanistically, IclRs repress transcription by two primary strategies: (1) directly blocking RNA polymerase binding or (2) interfering with transcription initiation by destabilizing interactions with the RNA polymerase α-subunit^41^. Despite their widespread presence, the functional diversity of IclRs remains underexplored, particularly in ecologically relevant species such as *P. putida*.

In this study, we demonstrate that LigR1 and LigR2 provide *P. putida* with a unique strategy to integrate both sugar import with aromatic compound metabolism, to repurpose shikimate intermediates for energy production. The LigR1 transcriptional regulator activates the *lig1* operon which participates in the import and processing of glucose and shikimate-derived compounds. LigR2 activates *ligR1* but also represses the *lig2* operon, which facilitates downstream utilization of aromatics for energy production. The repressive role of LigR2 is inactivated in response to 4-hydroxybenzoate, quinate, and protocatechuate accumulation. Dysregulation of LigR1 or LigR2 expression in *P. putida* KT2440 causes growth suppression on both glucose and aromatic substrates due to disruption in *lig1* and *lig2* operon expression. Deletion of *ligR1* also affects the morphology of *P. putida* KT2440 cells when grown in conditions supplemented with either glucose or 4-hydroxybenzoate. We put forth a model in which the *lig1* operon enables the import of glucose and shikimate-derived compounds and also the conversion of shikimate/quinate to dehydroshikimate (DHS). While the *lig2* operon expresses genes for the conversion of DHS to protocatechuate, which is cleaved to 2-pyrone-4,6-dicarboxylate by the actions of DSD and LigC, and funneled the resulting product into the protocatechuate meta cleavage pathway and eventually the TCA cycle for energy production^45,46^.

The coordinated regulation by LigR1 and LigR2 highlights the organism’s extraordinary metabolic flexibility and may explain its ability to proliferate in a highly competitive environment such as the rhizosphere. These findings offer new insight into microbial carbon metabolism and provide opportunities for engineering *P. putida* as a platform to develop biosensors or for the bioproduction of shikimate-derived compounds.

## Materials and Methods

### Bacterial strains, plasmids, media, and growth conditions

Bacterial strains and plasmids are listed in Supplementary Table 1, while primers are listed in Supplementary Table 2. *Escherichia coli* strains (DH5a plasmid host) and BL21 (protein production) were grown in Luria Bertani (LB) broth/agar media at 37°C. For *E. coli* strains carrying the pET28MOD or pBBR1-MCS2 plasmid, a concentration of 50µg ml^−1^ kanamycin was used. *E. coli* stains carrying the pEX18Ap plasmid were selected using 100μg ml^−1^ of ampicillin. *E. coli* stains carrying the pUCp20g or pmp220 plasmid were selected using 15μg ml^−1^ gentamicin and 10μg ml^−1^ tetracycline. *Pseudomonas putida* KT2440 was grown in LB broth/agar media or M9 minimal media supplemented with 25mM glucose, 4-hydroxybenzoate, quinate, and protocatechuate pH 7.4 at 30°C. *P. putida* KT2440 strains harbouring the pEX18Ap vector were selected with 500μg ml^−1^ carbenicillin or 30 µg ml^−1^ gentamycin. *P. putida* KT2440 strains carrying the pmp220 lacZ reporter plasmid or the pBBFLP recombinase plasmid were selected on 10µg ml^−1^ of tetracycline. *P. putida* KT2440 strains harbouring pBBR1-MCS2 or pUCp20g vector variants were selected with 50µg ml^−1^ kanamycin or 15µg ml^−1^ gentamicin, respectively. The *ΔligR2 P. putida* KT2440 strain was selected using 50µg ml^−1^ kanamycin and 30µg ml^−1^ chloramphenicol.

### Protein expression and purification

BL21 E. coli cells transformed with pET28MOD:*ligR1* and pET28MOD:*ligR2* were inoculated in LB with 50µg ml^−1^ kanamycin until the cultures reached the logarithmic phase (OD600nm 0.6 to 0.8). Protein expression was induced with 0.4mM isopropyl-*β*-thiogalactopyranoside (IPTG) and cultures were incubated at 16℃ for 16hrs. Cells were centrifuged and the resulting pellet was resuspended in 35mL of binding buffer (50mM TRIS-HCl, 5mM imidazole, 5% glycerol, 500mM NaCl) with 1mM benzamidine and phenylmethylsulfonyl fluoride (PMSF) protease inhibitor. Cell lysis was performed by a pre-chilled French press and sonication. Cell debris was removed by centrifugation and the recombinant protein was purified by nickel-nitriloacetic acid (Ni-NTA) affinity chromatography as described here^47^. For LigR1 and LigR2 used in electromobility shift assays, the soluble fraction was first poured through an equilibrated anion exchange cellulose DE52 column to remove fragmented DNA bound to the protein. 1mM of EDTA and 0.33mM of dithiothreitol (DTT) were added to each elution fraction and the presence of protein was confirmed using a ¼ dilution of Bradford Reagent. Tobacco Etch Virus (TEV) protease was used to cleave the hexa-histidine tags during overnight dialysis (10mM TRIS-HCl pH 7.5, 250mM NaCl, 5% Glycerol, *β*-mercaptoethanol) at 4℃. A second Ni-NTA affinity chromatography column was used to purify the un-tagged protein. The protein was dialyzed overnight at 4℃ and subsequently, concentrated by ultrafiltration to a final volume of 1mL.

### Crystallization, data collection, and structural determination of LigR1

The full-length LigR1protein comprising residues 1-256 was crystallized using the hanging drop-vapour diffusion method at 25℃. The protein was crystallized under conditions with 0.1M Tris-HCl pH 8.0, 0.2M Li_2_SO_4_, 0.020M NH_4_Acetate, 8% w/v PEG3350, and Arabidopsis root exudate compounds. Drops contained 2uL of protein solution (10mg mL^−1^) and 2uL of crystallization solution. Diffraction data were collected on Synchrotron APS Beamline 17-ID (Chicago, Illinois) with a Dectris Pilatus 6M Pixel detector. Images were processed with XDS and the intensities were scaled and merged with Aimless, and reduced using MOSFLM from the CCP4 suites^48^. The structure of LigR1was solved by molecular replacement using the IclR structure from *Rhodococcus* sp. RHA1 (PDB: 2IA2) and modified using AutoBuild from the Phenix Suites^49^. The structure was completed through iterative cycles of refinement using Refine and Coot, also from the Phenix Suites. The placement of acetate, 4-hydroxybenzoate and *trans*-cinnamic acid (*t*-cinnamic acid) were completed through Ligand Fit from the Phenix Suites using a 0.5 ligand occupancy threshold and set search parameters to the available ligand density.

### Differential scanning fluorimetry

20µL reactions containing 50µM of LigR1 or LigR2, 10X SYPRO orange protein stain (Sigma), and HEPES buffer (100mM HEPES pH 7.5 and 150mM NaCl) were prepared as previously described^50^. A 4-hydroxybenzoate and *t*-cinnamic acid saturation curve was generated using a concentration gradient from 0 to 5mM for LigR1. For LigR2, quinate, protocatechuate, 4-hydroxybenzoate at pH of 7.5 were tested at a concentration of 0.25mM. Four-replicates for each reaction were prepared in a 96-well plate and the thermal shift was measured using a temperature gradient from 25℃ to 95℃ at a rate of 1℃ per minute. All reactions were monitored with the BioRad CFX Real-Time System C1000 Thermal Cycler. Fluorescent outputs were recorded at 0.4℃ increments using the fluorescein amidites (FAM) channel at an excitation of OD450-490nm and emission at OD560-580nm. The protein melting temperatures were determined from the ‘CFX96 Manager’ software and graphed using the GraphPad Prism 10.3.1© software. The equilibrium dissociation constant (K_D_) value for LigR1 with 4-hydroxybenzote and *t*-cinnamic acid was determined using a non-linear regression analysis with one-site total binding using the GraphPad Prism 10.3.1© software.

### Electromobility Shift Assays (EMSA)

EMSA was performed with the purified full-length DE52 and Ni-NTA purified LigR1 or LigR2 protein as previously described^44^. A 250bp biotinylated DNA probe was amplified from the intergenic region upstream the of *rifI* gene. This *lig1* intergenic region probe was amplified from *P. putida* KT2440 gDNA using the 5’ end biotin-labeled forward primer and the unlabeled reverse primer for the *lig1* (LigR1_IR-F/R) and *lig2* operons (LigR2_IR-F/R). The 20µL binding reactions consisted of a binding buffer without EDTA, 15ng labeled DNA probe, 25ng µL^−1^ poly (dl-dC) non-specific competitor DNA, full length LigR1 protein at a concentration range from 20nM to 960nM. For LigR2, the same binding buffer conditions were used except the protein concentration tested ranged from 25nM to 1µM. For EMSAs testing the effect of ligands, 50nM of LigR1 was used and a varying concentration of 4-hydroxybenzoate from 0-500nM. For LigR2, we used 50nM of protein and 750nM of quinate, 4-hydroxybenzoate, protocatechuate, and *t*-cinnamic acid (pH 7.5). The biotinylated DNA: protein reaction mixture was incubated at 25℃ for 30 minutes. Following incubation, samples were separated on a 5% non-denaturing polyacrylamide gel prepared with ice-cold 0.5X Tris-borate EDTA (TBE; pH 7.5) buffer. Using a positively charged nylon membrane, the DNA was transferred by a semi-dry electroblot at 380mA for 30 minutes in 0.5x TBE. Subsequently, the DNA was crosslinked to the membrane for 30 minutes using a 300nm UV light. Membranes were washed 3 times in TBS-T before a 30-minute incubation with streptavidin-HRP at 25℃ (Cell Signaling Technology, diluted to 1/5000 in 5% BSA-T). The membranes were then washed in TBS-T again and the DNA was detected with Bio-Rad Clarity ECL Western Blotting Substrate. For the R1M, R2M, and R1R2M mutant DNA motifs were custom purchased from IDTDNA with the desired substitution. The same biotin-labeled primers (LigR1_IR-F/R) were used to amplify the mutated promoter fragments. Following amplification, the experiment was conducted as previously described above with 50nM of LigR1 or LigR2. To determine the fraction of bound DNA, we first quantified the pixel intensity (PI) of each band at different protein concentrations using the mean gray value in ImageJ. Next, for each protein concentration, we summed the pixel intensities of the shifted band(s) (SA, SB, …) and divided this value by the total intensity (sum of bound and unbound (UB) DNA) using the following equation:

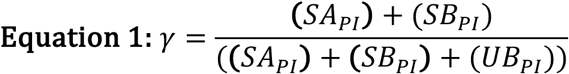

To calculate the transcription factor’s K_D_ for DNA binding, a non-linear regression analysis was conducted with one-site total binding using the GraphPad Prism 10.3.1© software.

### Gel Filtration

The native molecular weight of LigR1 was determined in the absence and the presence of 2.5mM 4-hydroxybenzoate (pH 7.5) at ambient temperature by a Pharmacia AKTA FPLC system with a UV-900 detector fitted with a Superdex 200 column (HiLoad 16/60, Pharmacia). LigR1was incubated at 25℃ with 2.5mM 4-hydroxybenzoate (pH 7.5) for 30 minutes before injection. Fast protein liquid chromatography was performed with mobile phases containing 50mM Tris-HCl, and 250mM NaCl (pH 7.5) at a flow rate of 0.5mL min^−1^ and injection volume of 100μL. Elution was monitored at OD280nm and identified fractions were further analyzed by SDS-PAGE to verify LigR1presence.

### Virtual ligand screening analysis of PP_2609

The structure-based virtual ligand modelling of LigR2 was performed using AutoDock Vina on the SwissDock interface^51–54^. The LigR2 AlphaFold PDB file and ligand structure using the advanced ligand search tool for quinate (ID: QIC), protocatechuate (ID: DHB), and 4-hydroxybenzoate (ID: PHB) were uploaded to the GUI^55^. To identify the location for docking, the LigR2 structure was superimposed to the LigR1 bound acetate structure in PyMOL to identify the approximate region of interest. A center grid box with dimensions x=6Å, y=-3Å, and z=-6Å and a box size of 12Å^3^ was generated to encompass the LigR2 binding site. Once the ligand of interest and the target protein was prepared, the sampling exhaustivity was set to 10 to increase the amount of computational effort to compute the binding affinity (kcal/mol). The target protein and the predicted ligand binding conformations were downloaded and viewed on PyMOL version 3.1.6.1.

### Generation of *ligR1* deletion in *P. putida* KT2440 by homologous recombination

To obtain a nonpolar markerless deletion of *ligR1* a two-step allelic exchange method with an engineered Gm resistant plasmid was used as previously described^56,57^. Two 300bp fragments upstream (SOEAB) and downstream (SOECD) of LigR1 were amplified from *P. putida* KT2440 gDNA using Phusion polymerase. Using the same polymerase, an overlap extension PCR was conducted to stitch the SOECD fragment to an FRT-flanked Gm cassette amplified from pS856. Subsequently, a second round of overlap extension PCR was conducted to stitch SOEAB to Gm:SOECD to produce a 1611bp SOEAB:Gm:SOECD fragment. Using the EcoRI/BamHI restriction enzymes, the SOEAB:Gm:SOECD fragment and the pEX18Ap plasmid were digested. Confirmed constructs were subsequently electroporated in *P. putida* KT2440. Carbenicillin (Cb) resistant colonies were plated on LB agar with 30µg mL^−1^ gentamicin (Gm30) and incubated overnight. The Gm resistant colonies were replica plated on LB agar Gm30 plates with 5% sucrose for sacB mediated counterselection overnight. Sucrose-resistant clones were streaked on LB-Gm30-5% sucrose and LB agar Cb500 plates to confirm for the loss of Cb resistance. The FRT-Gm cassette was excised by transforming pBBFLP expressing flippase (FLP) recombinase and a *sacB* counterselection marker. The transformants were selected on Tc LB-agar plates. To confirm the loss of Gm, Tc-resistant colonies were replica plated on LB-Gm30 agar plates to select for mutants unable to grow on Gm. The pBBFLP vector was cured from the unmarked *ΔligR1* mutant using sucrose counterselection.

### Complementation of *ΔligR1* and *ΔligR2* deletion and overexpression of LigR1 and LigR2

To complement the *ΔligR1* deletion, the WT *ligR1* gene was PCR amplified using the LigR1_Comp-F and LigR1_Comp-R primers and cloned into the pBBR1-MCS2-NdeI vector using NdeI and BamHI cut sites to yield the desired pBBR1-MCS2-NdeI:*ligR1* vector in DH5*α*. Following sequencing, the resulting plasmid was electroporated into either WT or *ΔligR1 P. putida* KT2440. Transformed *P. putida* KT2440 colonies were selected on kanamycin plates and PCR confirmed for the desired insert. The same approach was used for LigR2 using the LigR2_OE-F and LigR2_OE-R primers, however this construct was only transformed into the WT *P. putida* KT2440. To complement the *ΔligR2* deletion, the WT *ligR2* gene was PCR amplified using the LigR2_OE-F and LigR2_OE-R primers and cloned into the pucp20g-NdeI vector using NdeI and BamHI cut sites to yield the desired pUCp20g-NdeI:*ligR1* vector in DH5*α*. The resulting vector was electroporated into *P. putida* KT2440 *ΔligR2* and selected using kanamycin, chloramphenicol, and gentamicin plates.

### β-galactosidase reporter assay

Overnight *P. putida* KT2440 cultures grown in LB media harbouring the pmp220 vectors were pelleted and washed three times with M9 minimal media with 25mM glucose. The washed cells were diluted into fresh M9 minimal media with 25mM glucose at an OD600nm of 0.2 in 250mL flasks with respective antibiotics. β-galactosidase reporter strains also harbouring either the pBBR1-MCS2-NdeI or the pUCp20g-NdeI vector were supplemented with 0.4mM IPTG. The flasks were then placed in a 30℃ shaker at 200RPM until the OD600nm reached 0.6. Subsequently, the culture was split in 2.5mL aliquots into a 24-well round bottom plate and continued to grow for 2hrs in glucose with 200µM of inducer for 2hrs. Following induction, the final OD600nm of the culture was measured, while 1mL of the culture was pelleted and washed with 25mM Tris-HCl pH 7.5. The cell pellets were resuspended in Z-buffer [(60mM Na_2_HPO_4_, 40mM NaH_2_PO_4_, 10mM KCl, 1mM MgSO_4_, 20mM β-mercaptoethanol (pH 7.0)] with 3mg mL^−1^ lysozyme and incubated at 37°C for 1hr. Ortho-nitrophenyl-β-galactoside (ONPG) was added to a final concentration of 0.67mg mL^−1^, and all reactions were stopped after 15-minutes by adding 0.3M Na_2_CO_3_. Reaction start and stop times were recorded and centrifuged to pellet debris, and the OD420nm and OD550nm for each reaction was determined with a Cary 50 BioUV–Vis spectrophotometer. Miller units (MUs) were calculated using the following equation^58^:

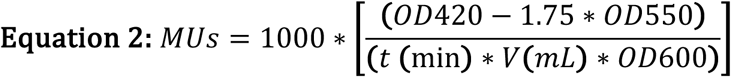

### Growth analysis of *P. putida*

Bacterial growth curves were performed by using *P. putida* growth cultures that were grown in LB media, pelleted, and washed three times with M9 salts. The washed cells were diluted into fresh M9 minimal media with 25mM glucose, 4-hydroxybenzoate, quinate, or protocatechuate (pH 7.4) to a final OD600nm of ∼0.08-0.1 in 250mL flasks in triplicates. The flasks were then placed in a 30℃ shaker at 200RPM for 24hrs. Bacterial growth was quantified by measuring the OD600nm of 1mL samples from the flasks every 3hrs for 9hrs, and then once at the 24hrs mark with a Cary 50 BioUV–Vis spectrophotometer.

### Cell lysis assay

Following 24hrs of growth, the final cell density of the cultures was measured at OD600nm. Subsequently, 1mL of culture was spun down at 5 x1000g for 10 minutes to pellet the cells. 300μL of the remaining growth media was added to a ¼ diluted Bio-Rad Protein Assay Dye Reagent to determine the total protein levels as a proxy for cell-lysis. The Cary UV spectrophotometer was blanked with ¼ Bio-Rad Protein assay with 300μL of sterile M9 minimal media with 25mM glucose. Protein content was measured at OD595nm in triplicates. The OD595nm value was divided by the OD600nm of the culture to normalize the value relative to the density of the culture.

### Crystal violet biofilm assays

Overnight cultures were diluted in fresh M9 minimal media with LB, 25mM glucose or 25mM 4-hydroxybenzoate to an OD600nm of 0.15 to a final volume of 20mL. The cells were poured into sterile Petri dishes containing 2 square glass coverslips that were placed inside. The cells were then placed in a 30℃ incubator not shaking for 24hrs. After 24hrs, the glass coverslips were removed from the petri dish. Using a Pasteur pipette, coverslips were washed three times with 0.85% sterile saline. Afterward, the coverslip was dried briefly from the backside using a Bunsen burner before staining the cells with 500μL of 0.1% crystal violet for 1 min. The excess crystal violet was washed three times with 0.85% saline and the coverslips were left to dry. Images were obtained using the Leica TCS SP5 Confocal Laser Scanning Microscope at 100x using oil immersion. Images were taken at 1x and 4x and zoomed in within a region of interest (ROI). Images were analyzed on Fiji (ImageJ) to obtain the bacterial cell lengths (μm)^59^. To determine statistical differences between the samples, a one-way ANOVA with Tukey’s multiple comparison test was conducted against the WT strain.

### Bioinformatic analyses

#### Multiple sequence alignment

The full-length amino acid sequences belonging to the LigR1, LigR2, PcaR, PobR, and IclR families were aligned through the Clustal Omega program (Supplementary Table 5)^60^. The alignment was annotated to show the ligand binding residues and their corresponding positions for each distinct class of IclRs. The full-length amino acid sequences for PobR (Q43992.1), PcaR (AAN66998.1), and IclR (P16528.1) were obtained from NCBI. The corresponding sequences were used as the query sequence in the Basic Local Alignment Search Tool (BLAST) to obtain the top 1000 hits (Altschul *et al.*, 1990; Gish and States, 1993)^61,62^. Subsequently, 3 unique species and the initial query sequence were used for alignment for each IclR.

#### Phylogenetic analyses

IclRs with biological functions in metabolism were used to query for lists of IclRs. These IclRs include: PobRs, CatRs, IclRs, LigR1s, LigR2s, and HmgRs. The top 1000 hits using BLAST for each IclR groups were filtered for non-redundant sequences and included together for a multiple sequence alignment using Clustal Omega. 131 sequences were included in a maximum likelihood phylogenetic analysis using the FastTreeMP program from XSEDE^63^. The phylogenetic analysis included 1000 bootstraps using the Jones-Taylor-Thornton (JTT) model and a penalty score of 1. The output from FastTreeMP was processed using the ITOL software and labeled using Inkscape^64^.

## Results

### Operon organization of genes within the *lig1* and *lig2* cluster

The *lig1* operon was the focus of this study since it houses a shikimate dehydrogenase (SDH) homolog in combination of genes encoding proteins that are annotated to be involved in energy sensing and carbon compounds utilization to fulfill the energy requirements of the cell. Interestingly, these processes are unrelated to the role of the shikimate pathway in an organism. Although this SDH homolog, RifI, was identified more than 15 years ago, its biological function has not been conclusively ascribed^42,43^. Comparative analysis of this SDH homolog revealed that it is positioned within a cluster of genes encoding two FAD-binding oxidoreductases, a NIPSNAP energy sensor, an MFS transporter, and a divergently transcribed IclR-type transcriptional regulator, *pp_2609* (LigR1)^65^ (Figure 1). Notably, this cluster lacked associated shikimate pathway enzymes, in contrast to other SDH homologs. Immediately downstream, is a second operon, which we name *lig2*, containing another IclR-type regulator (*pp_2601;* LigR2), a putative sugar phosphate isomerase (*pp_2603*), and *ligC*, encoding 4-carboxy-2-hydroxy-muconate-6-semialdehyde dehydrogenase-an enzyme previously demonstrated to mediate protocatechuate cleavage via the 4-5 pathway in *Sphingomonas paucimobilis* SYK-6^45^. Based on the combination of the encoded proteins between the two operons, we proposed that collectively they function in the utilization of shikimate-derived compounds to support the energy demands of the organism especially under starvation condition. This is consistent with previous findings for glucose and homogentisate utilization in *P. putida* where the genes encoding the enzymes are organized modularly for distinct portions of the utilization pathway^7,19^. We thus proposed an interplay between both operons to prioritize carbon compounds usage based on their availability in the environment. This investigation provided a unique opportunity to challenge the established role of the shikimate pathway in microbes, focusing instead on the use of its intermediate compounds as alternative source for energy. In context of *P. putida’s* lifestyle within the plant rhizosphere, we hypothesize that additional catabolic routes could be beneficial in a dynamic and competitive environment for nutrients and space. To initiate our investigation, we focused on the transcriptional regulation of these two operons and conducted detailed biochemical and molecular studies both *in-vitro* and *in-vivo* to deconvolute their role.

**Figure 1.**
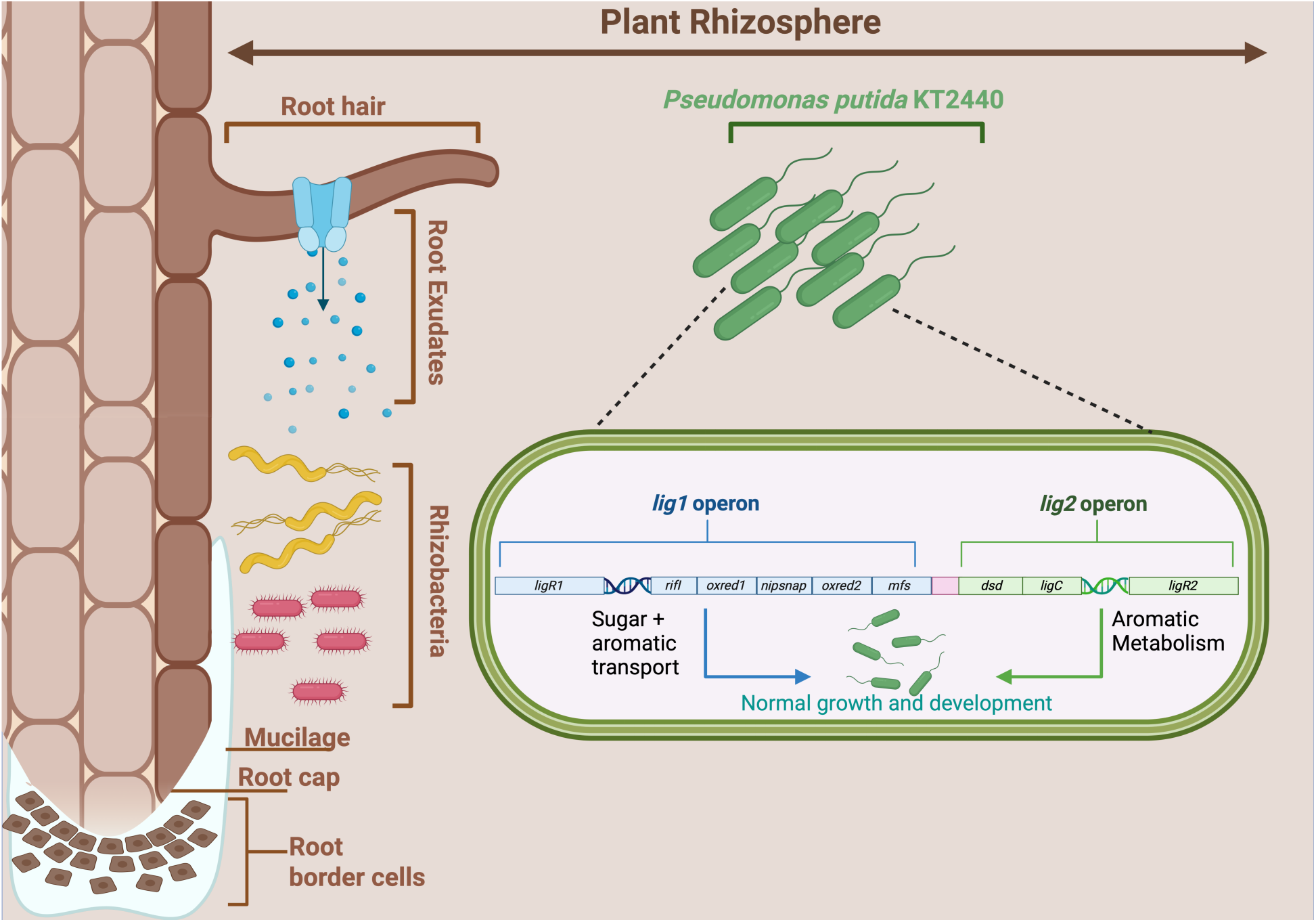
Genomic organization of the *Pseudomonas putida* KT2440 *lig1* and *lig2* operons. Illustration of the plant rhizosphere, which harbors a diverse microbial community, including *P. putida* KT2440 (depicted in green). Microorganisms within the rhizosphere compete for root exudate compounds such as sugars, organic acids, and aromatic molecules released from plant roots into the surrounding soil. Within the genome of *P. putida* KT2440, the *lig1* (blue) and *lig2* (green) operons contribute to the uptake and metabolism of aromatic compounds, supporting bacterial growth and development. The *lig1* operon (genes *pp_2608* to *pp_2604*) encodes a shikimate-like dehydrogenase homolog (*rifI*), two FAD-binding oxidoreductases (*oxred1* and *oxred2*), a 4-nitrophenylphosphate non-neuronal SNAP25-like protein (*nipsnap*), and a major facilitator superfamily (*mfs*) transporter. Upstream of *lig1* is a divergently transcribed IclR-family transcriptional regulator, *ligR1.* The *lig2* operon (genes *pp_2602* to *pp_2603*) encodes a putative dehydroshikimate dehydratase (*dsd*) and a 4-carboxy-2-hydroxy-6-semialdehyde dehydrogenase (*ligC*), and a second IclR-family transcription factor, *ligR2*. The two operons are separated by approximately 150 base pairs, indicated by the pink box between them.

### LigR1 binds to lig1 promoter, while LigR2 binds both lig1 and the *lig2* promoters

To investigate the role of the *lig1* and *lig2* operons in *P. putida* KT2440, we used electrophoretic mobility shift assays (EMSAs) to examine how the respective transcriptional regulators bind to their intergenic regions. For the *lig1* operon, a biotin-labeled probe spanning +23 bp (relative to the *ligR1* translation start site) to +72 bp (within the *rifI* gene) was used in EMSA studies (Figure 1Ai). The biotinylated probe was incubated with increasing concentrations of LigR1 (0–0.96µM). A control containing only the biotinylated probe (no protein) was included to represent unbound DNA (Lane 1). At 0.12µM LigR1, two distinct shifted bands (S1 and S2) appeared, indicating protein-DNA complex formation (Lane 4). At higher protein concentrations (>0.12µM), the intensity of the faster mobility band (S1) decreased, while a third, slower-migrating band (S3) emerged at 0.24µM LigR1 (Lane 5). This S3 band intensity further increased at the highest concentration of protein (0.96µM, Lane 7). Concurrently, the S2 band increased in intensity with increasing LigR1 concentrations (Lanes 4–7). The fraction of bound DNA was quantified and plotted against LigR1 concentration (Equation 1, Materials and Methods) (Figure 1Aii). At 0.96µM LigR1, the bound DNA fraction reached 0.95, yielding a K_D_ value of 0.22µM for LigR1 binding to the probe.

For the *lig2* operon, we designed a similar biotin-labeled probe spanning +48 bp (relative to the *ligR2* translation start site) to +18 bp (within the *ligC* gene). The biotinylated probe was incubated with increasing concentrations of LigR2 (0–1µM), with a no-protein control (Lane 1) representing unbound DNA (Figure 1Bi). Protein-DNA complex formation was evident at LigR2 concentrations as low as 0.025µM, also yielding two distinct shifted bands (S1 and S2; Lane 2). The intensity of these bands increased with LigR2 concentrations up to 0.125µM. At 0.25µM LigR2, the majority of complexes migrated as the S2 band, accompanied by the emergence of a third shifted species (S3). The S3 band intensity rose progressively, reaching its maximum at 1µM LigR2 (Figure 1Bi). Quantification of bound DNA revealed a saturation binding fraction of 0.97 at the highest LigR2 concentration (1µM), with a dissociation constant K_D_ of 0.15µM (Figure 1Bii).

We examined whether LigR1 could bind to the biotinylated *lig2* intergenic region using the similar EMSA assay. Increasing LigR1 concentrations (0–1µM) did not induce a mobility shift in the *lig2* DNA probe, indicating a lack of binding (Supplementary Figure 1Ai-ii). In contrast, incubation of LigR2 with the biotinylated *lig1* probe resulted in a single shifted band, absent in the no-protein control (Lane 1) (Figure 1Ci). This shift was detectable at LigR2 concentrations as low as 0.05µM and reached a maximal intensity by 0.5µM. Quantitative analysis revealed that the bound *lig1* fraction saturated at 0.65µM, with residual unbound DNA remaining at the lowest observed concentrations (Lanes 6–8) (Figure 1Cii). The dissociation constant K_D_ was 21.65µM. As an additional control, competition assay was conducted using increasing concentrations of unlabelled probe to confirm specificity of the interactions between LigR1 and LigR2 with their respective promoter regions (Supplementary Figure 1B-E). Together, these findings demonstrate that while both LigR1 and LigR2 bind their cognate intergenic regions, only LigR2 exhibits cross-binding to *lig1* intergenic region.

### LigR1 and LigR2 DNA-binding occurs at a specific recognition TAAT motifs

Systematic analysis of the DNA-binding signatures of 1,340 IclR-family transcription factors have been classified by their recognition motifs into two distinct groups: A/T-rich palindromic sequences (Group 1) and the consensus GTNCG-N5–6-CGNAC motif (Group 2)^66^. Our analysis identified two putative operator regions in the *lig1* promoter (R1: ATCATGAC**T**G**CGATTA**T**CG**CA; R2: ACCAGC**CGATTA**CTG**CG**AT) that closely matched the Group 2 architecture (Supplementary Figure 2Ai). To functionally validate these sites, we generated targeted mutations by converting the conserved ATTA sequences to GCTG in R1 (R1M), R2 (R2M), and a double mutant (R1R2M) in the *lig1* promoter region (Supplementary Figure 2Ai). EMSA demonstrated complete complex formation (S2/S3) with wild-type *lig1* intergenic region with the LigR1 transcription factor (Lane 2), while all mutated probes showed partial binding disruption of LigR1, as demonstrated by loss of the S3 band. This binding disruption was most prominent with the R1R2M double mutation (Lanes 4,6,8) (Supplementary Figure 2Aii). Quantification of the DNA-bound fraction revealed approximately 50% and 37% reduction in binding to the R1M and R2M probes, respectively (Supplementary Figure 2Aii). More strikingly, approximately 70% reduction in LigR1 binding affinity was observed for the R1R2 mutated probe. These results demonstrate that both R1 and R2 ATTA sites in the *lig1* promoter region facilitate high-affinity LigR1 binding.

We next examined how *lig1* intergenic mutations affect LigR2 binding. Sequence alignment of the *lig2* intergenic region with *lig1* R1M, R2M, and R1R2M mutants revealed conservation of the ATTA motif at the R1 position (Supplementary Figure 2Bi, 2C), while the R2 site contained a divergent CTTC sequence. EMSA analysis of GCTG-substituted *lig1* sequences (Supplementary Figure 2Bii) showed distinct binding patterns: the WT *lig1* intergenic region produced a single shifted band (S1) with residual unbound probe (Lane 2), whereas the R1M mutant exhibited an additional shifted band (S2) with no free probe (Lane 4). The R2M mutant displayed only the S1 band with reduced free probe intensity (Lane 6), while the R1R2M double mutant displayed both S1 and S2 bands with partial free probe persistence (Lane 8). Quantitative analysis demonstrated a 36.5% increase in bound DNA fraction for R1M, indicating that LigR2 binding is mediated primarily through the ATTA motifs at the R1 and R2 position. Overall, these data suggests that LigR1 and LigR2 share a common ATTA binding motif.

**Figure 2.**
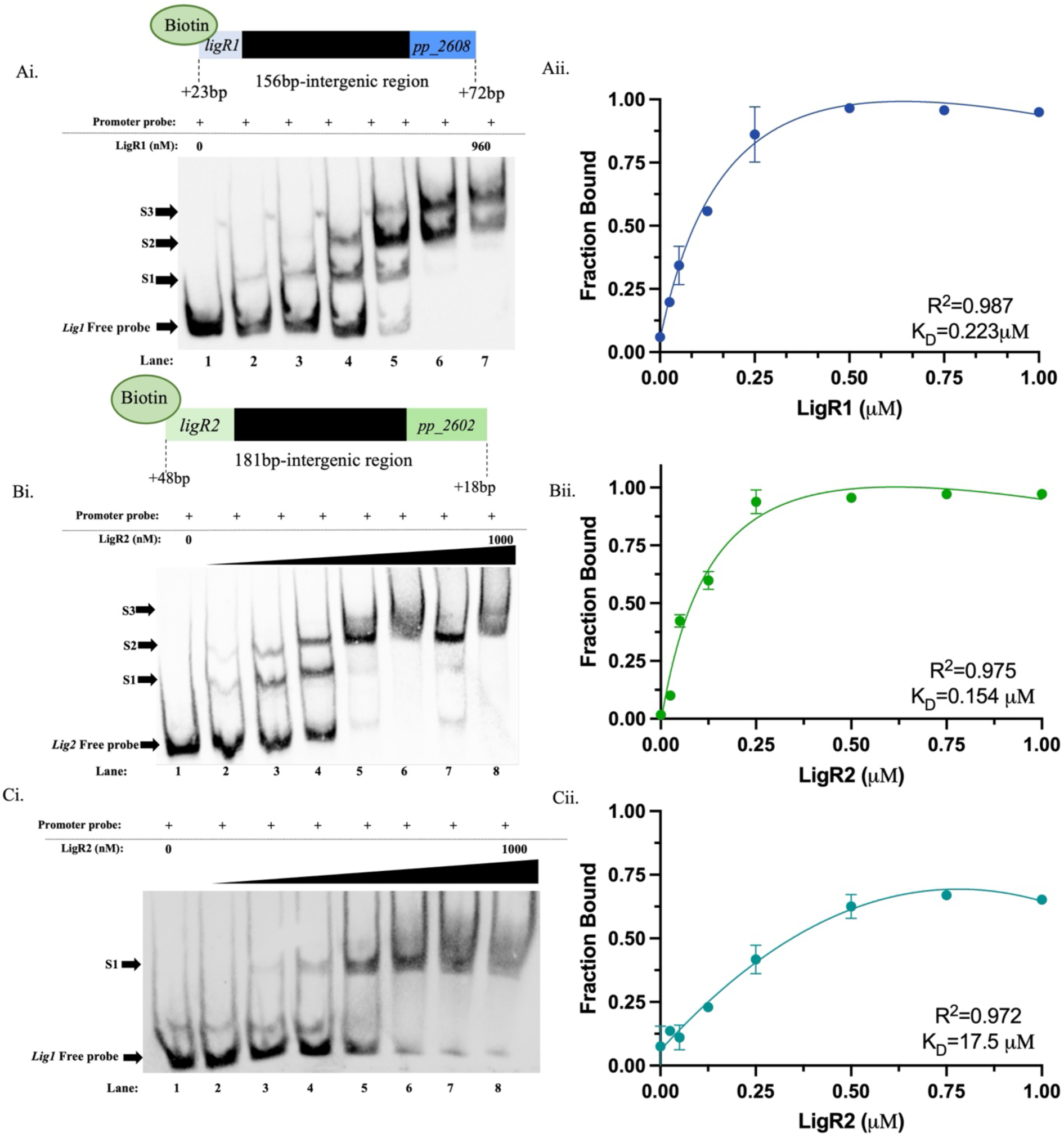
EMSA analyses of LigR1 and LigR2 binding to the *lig1* and *lig2* intergenic regions. Ai) Schematic of the 251bp biotinylated *lig1* DNA probe, encompassing the intergenic region between *ligR1* (+23bp relative to the translation start site) and *rifI* (+72bp). A biotin label was added to the 5’ end for detection using streptavidin-HRP for all DNA probes. Below the schematic, the EMSA gel investigated concentration-dependent binding of LigR1 to the *lig1* promoter region. Lanes 1-7 contain increasing concentrations of LigR1 (0, 20, 60, 120, 240, 480, and 960nM). Mobility shift positions are labeled as Shift 1 (S1), Shift 2 (S2), and Shift 3 (S3). Aii) Quantification of *lig1* promoter binding by LigR1, shown as the fraction of DNA bound at each concentration, based on band intensity comparisons between the bound and unbound DNA. Bi) Schematic of the 247bp biotinylated *lig2* DNA probe, encompassing the intergenic region between *ligR2* (+48bp relative to the translation start site) and *ligC* (+18bp). Below the schematic, the EMSA gel investigated concentration-dependent binding of LigR2 to the *lig2* promoter region. Lanes 1-8 contain increasing concentrations of LigR2 (0, 25, 50, 125, 250, 500, 750, and 1000nM). Bii) Quantification of *lig2* promoter binding by LigR2, shown as the fraction of DNA bound at each concentration, based on band intensity comparisons between the bound and unbound DNA. Ci) EMSA analysis of LigR2 binding to the *lig1* promoter probe. Lanes 1-8 contain increasing concentrations of LigR2 (0, 25, 50, 125, 250, 500, 750, and 1000nM). Cii) Quantification of *lig1* promoter binding by LigR2, shown as the fraction of DNA bound at each concentration, based on band intensity comparisons between the bound and unbound DNA. All quantification data were fitted to the non-linear equation: Y = Bmax/(K_D_ + X) + NSX, where Bmax is the maximum fraction bound and NS the slope of the non-linear component, representing non-specific binding. The coefficients of determination (R^2^) and K_D_ values obtained are shown in the graphs for each graph. EMSA assays were performed in duplicate. Error bars indicate the Standard Error of the Mean (SEM). Concentration of probe used in EMSA assays was 15 ng uL^−1^ as described in the Materials and methods.

### The LigR1 and LigR2 structures resemble IclR-type transcriptional regulators

To investigate the molecular function of LigR1, we determined its crystal structure using Molecular Replacement with a previously determined IclR structure from *Rhodococcus* sp. RHA1 (PDB: 2IA2) as a template model. The final model was refined to 2.11Å resolution and contained four polypeptide molecules within the asymmetric unit (Supplementary Table 3 for data collection and refinement statistics, PDB: 9E6A). For the LigR1_A protomer, the structure encompassed residues Ile6 to Val254. For the B protomer, the structure contained residues from His7 to Asp256. For protomer C the structure encompassed residues Pro4 to Leu255. A single protomer of LigR1 contained an N-terminal helix-turn-helix DNA binding domain (DBD) and a C-terminal effector binding domain (EBD) (Figure 3A). The DBD is connected to the EBD α-helical linker followed by a short loop spanning 3 residues (Ser84 to Glu86) (Figure 3A). The C-terminal domain constitutes 6 α-helices and 6 anti-parallel *β*-sheets, which organize into a GAF-like domain superfamily (Figure 3B)^37^. Within the EBDs, we observed electron density for acetate from the crystallization condition in all the protomer chains, encompassed by the *β*-sheet. Two protomers (Chain A&C, Chain B&D) dimerized at the DBD through interactions at the dimerization domains α2 with α2’, α2 with α4’, and α5 with α5’ (Figure 3C). Both dimers assembled into tetramers with the DBDs facing outwards and the EBDs occupying the center of the protein structure.

**Figure 3.**
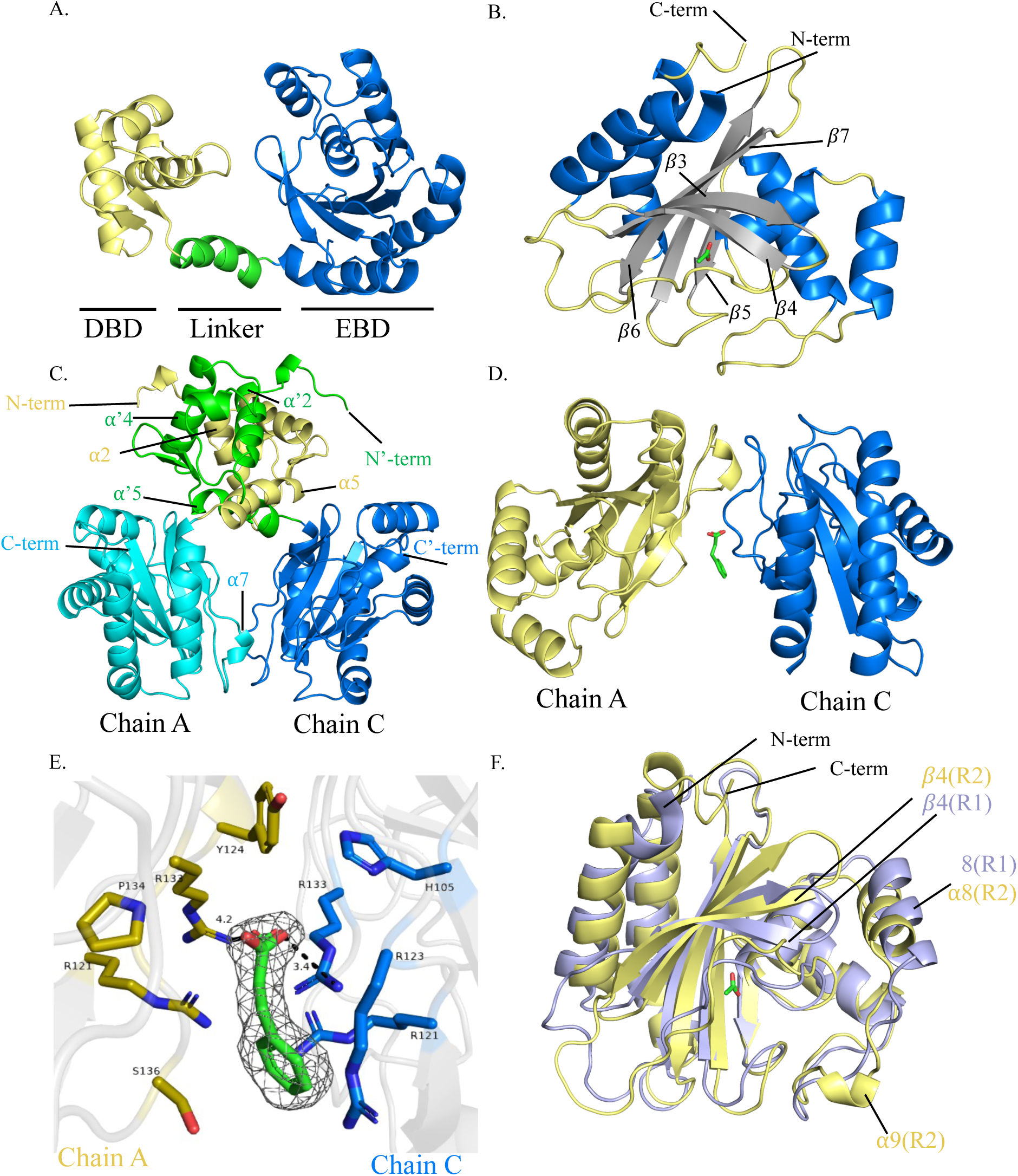
Overview of the LigR1 crystal structure and LigR2 AlphaFold model. A) A Ribbon representation of the LigR1 monomer (PDB ID: 9E6A). The helix-turn-helix (HTH) DNA binding domain (DBD) is shown in yellow, the ɑ-helical linker in green, and the effector binding domain (EBD) in marine blue. B) Ribbon representation of the EBD from protomer D complexed with acetate (green). Secondary structural elements are coloured-coded as follows: loops (yellow), *β*-sheets (grey), and ɑ-helices (marine blue). The N- and C-termini and the *β*-strands forming the anti-parallel sheet are annotated. C) Dimerization interface between LigR1 protomers at their DBDs. Chain A (yellow) and Chain C (green) DBDs interact at the ɑ2, ɑ4, ɑ5 helices of the HTH motif. Additional contact occurs at the ɑ7 helix of Chain A’s EBD (cyan) and Chain C’s EBD loop (marine). The N- and C-termini of both protomers are labelled. The prime (’) notation indicates elements from Chain C. D) Ribbon diagram showing Chain A (yellow) and C (marine) EBDs docked with *t*-cinnamic acid (green). E) Close up view of the *t*-cinnamic acid (green) binding site between EBD residues of Chain A (yellow) and Chain C (marine). Residue side chains are shown using single-letter amino acid codes. The dark grey mesh represents the electron density corresponding to *t*-cinnamic acid, as resolved in Phenix. F) Structural superimposition of LigR1 Chain D’s EBD (purple), bound to acetate (green) with the AlphaFold predicted model of LigR2 (yellow) EBD. Differences in secondary structure are highlighted in their respective colours. All images were generated from PyMOL version 3.1.6.1

Additional electron density was observed at the dimer interface and is within proximity to a sulfate ion. The electron density is bulky and likely represent a bound aromatic compound since the crystallization condition was supplemented with *A. thaliana* root exudates, which is highly represented by monolignol-type compounds. We therefore attempted to fit monolignols-type compounds, p-coumaric acid, p-coumaryl alcohol, ferulic acid, caffeic acid, and *t*-cinnamic acid within this density. Of the ligands that were docked using the Ligand Fit program on Phenix, *t*-cinnamic acid had the highest coefficient correlation of 0.67 with a ligand occupancy of 0.5 (Figure 3D). The binding site analysis identified two arginine residues from Chain A (R121, R133) and four arginine residues from Chain C (R121, R123, and R133) that are within close proximity to the docked *t*-cinnamic acid. R133 from chain A and C formed hydrogen bonds with the carboxylate group of *t*-cinnamic acid (Figure 3E). To date, no other IclRs have been reported to have a secondary binding site, providing a potentially novel mechanism of regulation through allosteric control.

We compared the AlphaFold-predicted LigR2 structure with the LigR1 crystal structure to identify potential structural differences. Similar to LigR1, LigR2 exhibited conserved IclR-type features including a helix-turn-helix DBD connected via an α-helical linker with a short loop spanning a single G79 residue, to the EBD. The EBD architecture also consisted of a 6-stranded anti-parallel *β*-sheet representative of the GAF-like domain (Figure 3F). Structural alignment revealed three notable differences between LigR2 and Chain D of LigR1. First, the *β*3-*β*4 loop adopted an upward orientation in LigR2 compared to the downward projection in LigR1. Second, the α8 helices of the two structures were orientated perpendicularly relative to each other. Third, LigR2 contained an additional α9 helix, but lacked the α7 helix seen in LigR1’s structure. These differences between LigR1 and LigR2 occurred despite the strong overall conservation, with an RMSD of 2.814 Å across aligned residues in PyMOL. The structural variations seen particularly in the EBD, provide evidence of either the confirmational changes occurring during ligand binding or the potential functional divergence between these homologous regulators.

### LigR1 and LigR2 are evolutionarily distinct and share minimal sequence conservation

To investigate the relationship between LigR1 and LigR2 with other characterized IclR-type transcriptional regulators, phylogenetic analysis was conducted together with groups of functionally distinct IclRs with high sequence similarity. These IclRs include: IclR which regulates acetate utilization, HmgR which controls homogentisate metabolism, PcaR which activates protocatechuate metabolism through the *β*-ketoadipate pathway, PobR which activates 4-hydroxybenzoic acid metabolism, and CatR required for the catabolism of catechol^20,30,31,39^. A maximum likelihood analysis of these IclRs produced seven major clusters supported with high bootstrap values, exceeding 0.96 (Figure 4A). This analysis positioned LigR1 as a distinct but closely related lineage to PobR, implying functional parallels in regulation of aromatic compounds metabolism. Conversely, LigR2 clustered independently from the characterized IclRs, suggesting that the role of this transcriptional regulator is either functionally broad in terms of the ligands it responds to or is highly specialized and differs from the other IclRs. Multiple sequence alignment was conducted between LigR1 and LigR2 with representative species from the PobR, PcaR, and IclR clusters to investigate conservation and the physiochemical properties of the ligand binding residues (Figure 4B, Supplementary Table 4, Supplementary Figure 3). Of the nine ligand binding residues of PcaR, only two residues are conserved in LigR1 and LigR2. For PobR, only one of the seven ligand binding residues is conserved in LigR1 and none in LigR2. Analysis with IclR did not identify any conserved ligand binding residue with LigR2. Interestingly, these proteins all show a conserved ligand binding pocket with similar physiochemical properties, the level of conserved ligand binding residues is quite variable.

**Figure 4.**
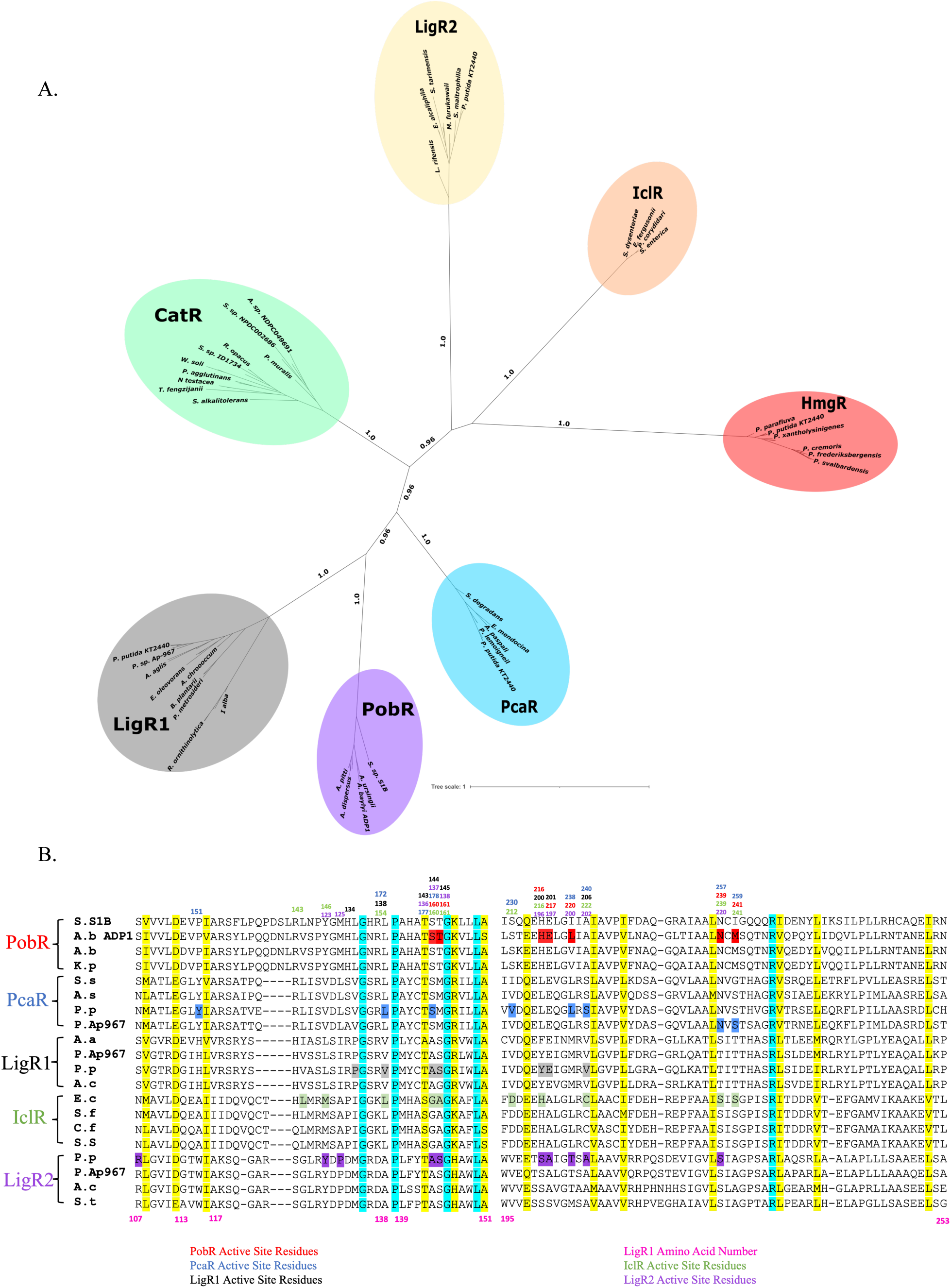
Phylogenetic analysis and sequence alignment of LigR1 and LigR2 with characterized IclR-type transcription factors. A) NCBI BLAST sequence database was used to query gene homologs for LigR1, LigR2, CatR, IclR, PcaR, PobR, and HmgR. The sequences were aligned using Clustal Omega and phylogenetic analysis was conducted by Fast Maximum Likelihood (FastTreeMP) on XSEDE using the Jones-Taylor-Thorton (JTT) model with 1000 bootstraps. The final tree was displayed using iTOL with annotations drawn using Inkscape. Major nodes supported by ≥80% bootstrap scores are indicated on the tree. B) Multiple sequence alignment for PobR, PcaR, IclR, LigR1 and LigR2 groups from select species were aligned using Clustal Omega. Turquoise-coloured residues show sequence conservation, while the yellow-coloured residues show similar amino acid. Active site residues for the corresponding IclR transcription factors are coloured in red (PobR), blue (PcaR), black (LigR1), green (IclR) and purple (LigR2) with the corresponding amino acid number above the alignment.

### Gene expression analysis by *β*-galactosidase reporter assay

To elucidate the regulatory roles of LigR1 and LigR2, we performed systematic gene expression analyses in *P. putida* KT2440 using a *β*-galactosidase reporter assay. Reporter constructs were designed to evaluate the role of LigR1 and LigR2 on both *lig1* and *lig2* operon expression and also on autoregulation (Figure 5). To identify the metabolites that could activate *lacZ* expression, we generated three independent chemical mixtures containing five to nine compounds based on their biological role in the cell (Supplementary Figure 4A). The three groups of mixtures can be classified as sugars, aromatic compounds, and intermediates of the TCA cycle. Only the mixture containing aromatic compounds: protocatechuate, *γ*-butyric acid, 4-hydroxybenzoate, *t*-cinnamic acid, quinate, shikimate, leucine, and phenylalanine activate *lacZ* expression in context of *lig2* operon expression (Supplementary Figure 4B). When tested individually, three compounds, protocatechuate, quinate and 4-hydroxybenzoate stimulated LacZ activity with quinate showing the strongest effect (702 MU) on *lig2* operon expression compared to glucose (94 MUs) (p<0.0001) (Figure 5B). For *ligR2* autoregulation, we did not observe strong *lacZ* activity with any mixture tested (Figure S4A). We only observed a moderate increase in in LacZ activity with quinate (248MUs, p=0.0153), protocatechuate (257 MUs, p=0.005), and 4-hydroxybenzoate (274 MUs, p=0.0007) compared to glucose (183 MUs). We also investigated the consequences of overexpressing LigR1 and LigR2 on *lig2* operon expression. In both cases, overexpressing LigR1 and LigR2 reduced *lig2* operon expression similar to the glucose control condition even with aromatic compounds supplementation. To rule out artifactual effect from the plasmid, we tested *lig2* operon expression with the empty plasmid and did not observe a repressive effect on *lacZ* expression. This indicated a coordinated effect of LigR1 and LigR2 on repressing *lig2* operon expression.

**Figure 5.**
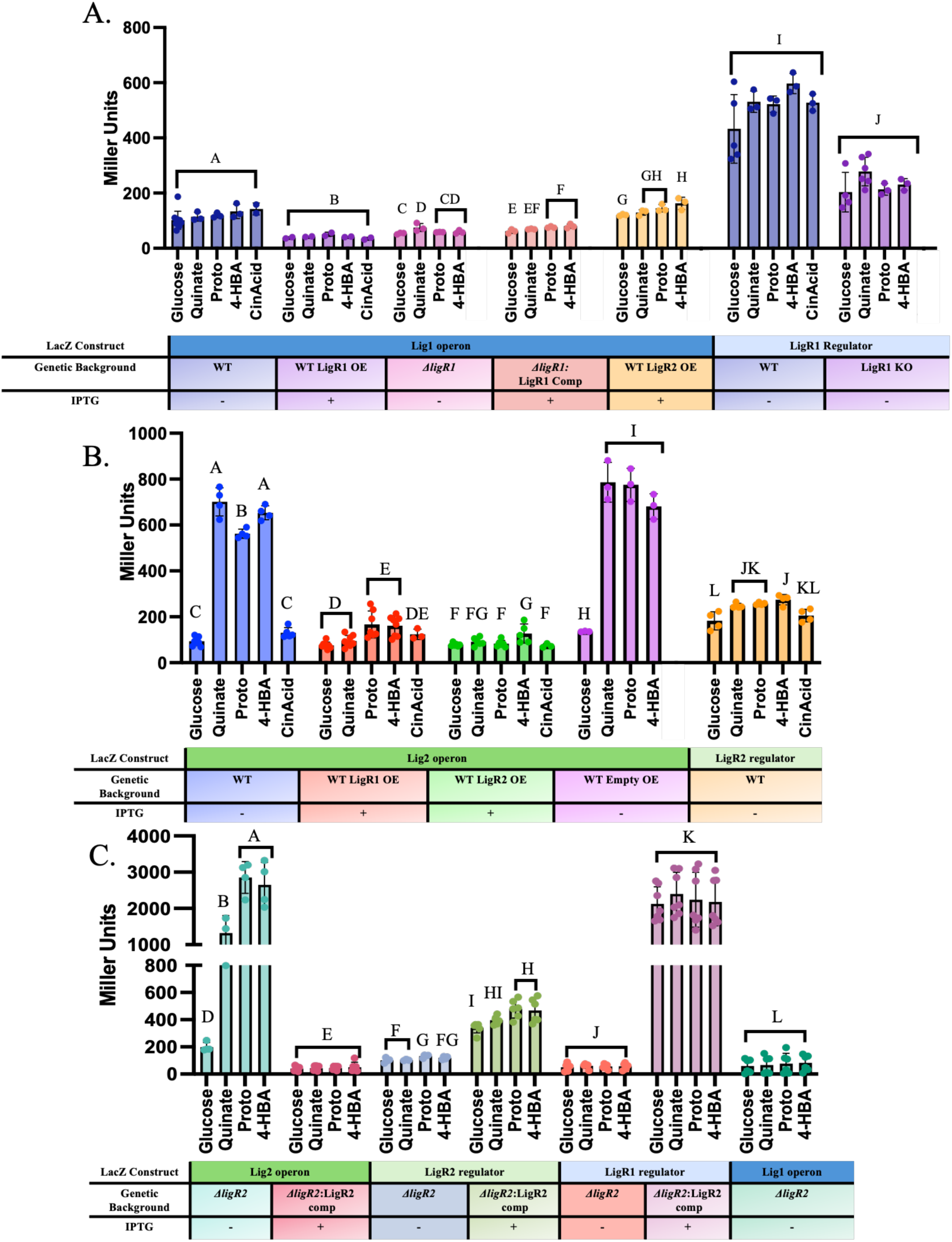
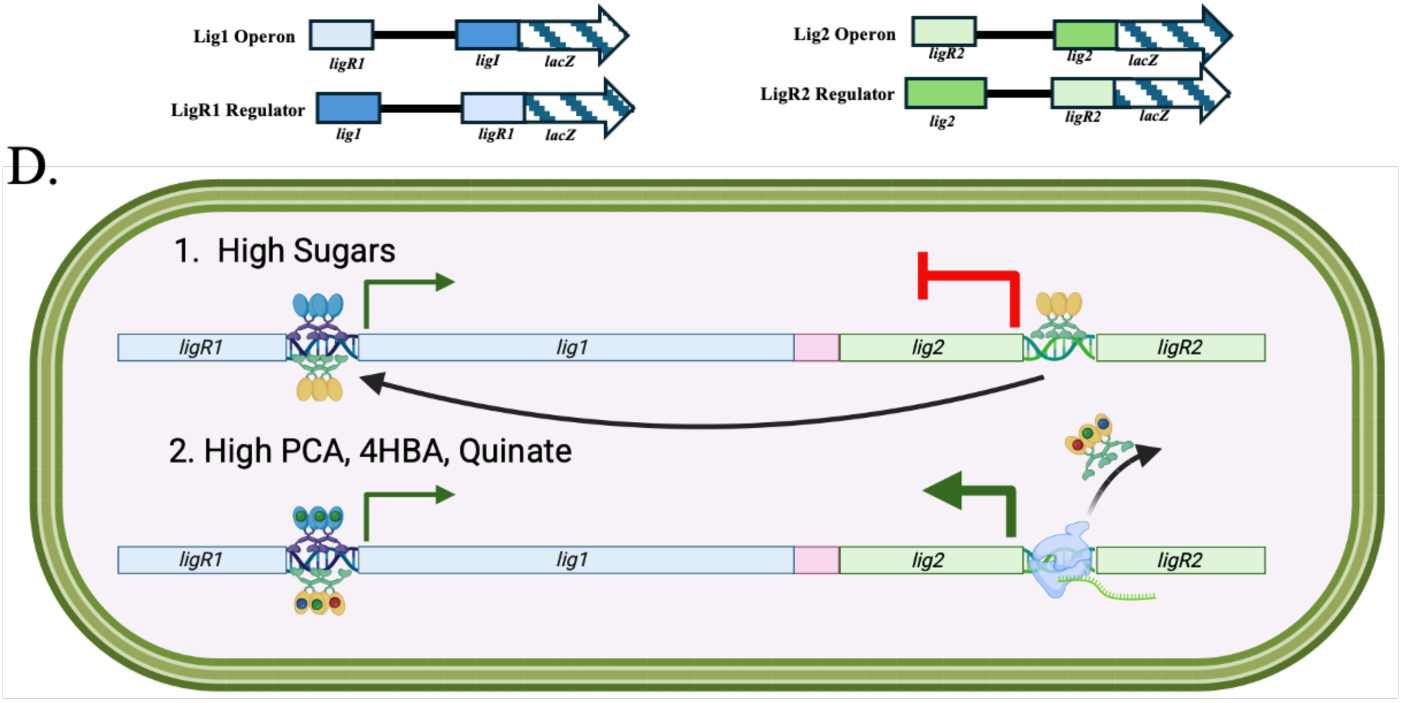
Gene expression analysis of LigR1 and LigR2 using a β-galactosidase reporter assay. A) Gene expression analysis of the *lig1* operon and autoregulation of the *ligR1* regulator using the 251 bp *lig1* intergenic region cloned upstream of a *lacZ* reporter gene. B) Gene expression analysis of the *lig2* operon and autoregulation of the *ligR2* regulator using the 247 bp *lig2* intergenic region cloned upstream of a *lacZ* reporter gene. C) Gene expression analysis of the *lig1* and *lig2* operons, as well as *ligR1* and *ligR2* autoregulation, in the Δ*ligR2* and Δ*ligR2*:LigR2 complemented backgrounds. All investigations were conducted in either wild-type (WT), Δ*ligR1*, Δ*ligR1*:LigR1, Δ*ligR2*, or Δ*ligR2*:LigR2 *P. putida* KT2440. Gene expression was induced with 200µM of either quinate, 4-hydroxybenzoate, protocatechuate, or *t*-cinnamic acid, in the presence (+) or absence (–) of 0.4mM IPTG. Data points represent biological replicates. Statistical analysis was performed using one-way ANOVA with Tukey’s multiple comparisons test. Letters above the bars indicate compact letter displays, where different letters represent statistically significant differences at a 95% confidence interval (CI). Error bars indicate the SEM. D) Summary schematic of the β-galactosidase reporter assay under high sugars versus high aromatic compounds. Under high sugars, LigR2 and LigR1 bind to the *lig1* promoter to activate expression of the *lig1* operon. LigR2 binds to the *lig2* promoter and represses expression of the *lig2* operon. Under high aromatics, LigR2 and LigR1 bind to the *lig1* promoter to activate expression of the *lig1* operon. Quinate, protocatechuate, and 4-HBA binding to LigR2 inactivates repression enabling *lig2* operon expression.

Similarly, we investigated *lig1* operon and *ligR1* regulator expression using a similar LacZ reporter system. We did not observe a difference in *lig1* operon expression using the three chemical mixtures described above (Supplementary Figure 4A). We also tested the metabolites individually and also did not observe a difference in operon expression compared to the glucose control (101-131 MUs) (Figure 5A). In contrast to the operon expression, LacZ activity for the *ligR1* regulator expression exhibited a 4-fold increase with glucose, but the expression did not change with aromatic compound supplementation. Next, we tested the effects of ectopically overexpressing LigR1 and LigR2 by inducing their expression with IPTG. In all cases, *lig1* operon expression did not show a detectable change in *lacZ* reporter activity. Finally, we generated a *ligR1* gene knockout in *P. putida* (*ΔligR1*) to determine its effect on *lig1* operon and regulator expression. This *ΔligR1 P. putida* strain showed a marked decrease in both *lig1* operon (53-75MUs) and *ligR1* expression by ∼2.5 fold compared to the WT background (203-278 MUs). As a follow up experiment, we did not detect any lacZ activity with the ∼150bp region between the two operons (Supplementary Figure 4C).

We also conducted *β*-galactosidase reporter assay in the *P. putida ΔligR2* background to determine the effect of LigR2 on *lig1* and *lig2* operon expression (Figure 5C). For *lig2* operon expression we observed high LacZ activity using quinate (1322 MUs), protocatechuate (2859 MUs), or 4-hydroxybenzoate (2652 MUs) as the inducing agent, while the complementation strain showed reduced *lig2* operon expression with IPTG supplementation. In contrast we observed a decrease in LacZ activity associated with repression of *ligR2* regulator expression, with quinate (103 MUs), protocatechuate (131 MUs), or 4-hydroxybenzoate (118 MUs), which reverted to similar expression level to WT in the complementation strain with IPTG supplementation (329-429MUs). For *ligR1* regulator expression in the *ΔligR2* background we observed a striking decrease in LacZ activity to 50 MUs irrespective of which compound tested. Further, we were able to revert *ligR1* regulator expression in the LigR2 complemented strain with IPTG supplementation. Finally, for *lig1* operon expression we also observed a decrease in expression to 59-82 MUs for the tested compounds. Overall, we concluded that both LigR1 and LigR2 functions as an activator for the *lig1* operon, but as repressor for the *lig2* operon when there is high concentration of sugars but low concentration of aromatic compounds (Figure 5D). In contrast, when there is a high concentration of aromatic compounds, LigR2 repression of *lig2* operon is relieved to facilitate gene expression. The *lig1* operon continues to be expressed in both scenarios.

### LigR1 and LigR2 effector binding analysis

To elucidate how the identified metabolites from the *β*-galactosidase reporter assay influence protein biophysics, we performed differential scanning fluorimetry (DSF) and size-exclusion chromatography. DSF analysis showed that *t*-cinnamic acid and 4-hydroxybenzoate increased LigR1’s thermal stability by 2.0°C and 2.1°C, respectively (Figure 6A). The K_D_ value for *t*-cinnamic acid and 4-hydroxybenzoate binding was determined to be 2.54 x 10^−4^ M and 2.31 x 10^−4^ M, respectively. In contrast, LigR2 displayed thermal destabilization with 0.25mM protocatechuate, 4-hydroxybenzoate, and *t*-cinnamic acid (Figure 6B). Gel filtration chromatography revealed that 2.5mM 4-hydroxybenzoate induced subtle changes in LigR1’s oligomeric state (Supplementary Figure 6A,B). The *apo* protein primarily eluted as a 57.5 kDa dimer, while 4-hydroxybenzoate treated samples showed both a dimer peak at 57.5 kDa and a new tetramer peak at 115.2 kDa. We were unable to determine the effects of *t*-cinnamic acid on LigR1 and LigR2 oligomeric state due to the absorbance interference with the metabolite at 280nm. Quantitative analysis demonstrated the tetramer population increased ∼12-fold based on AUC values, from 3.98 in *apo* conditions to 48.78 with ligand, while the dimer population showed a modest 6% increase from 932 to 992.2 (Supplementary Figure 6B). These findings establish that 4-hydroxybenzoate interacts with LigR1, inducing measurable shifts in its oligomeric state.

**Figure 6.**
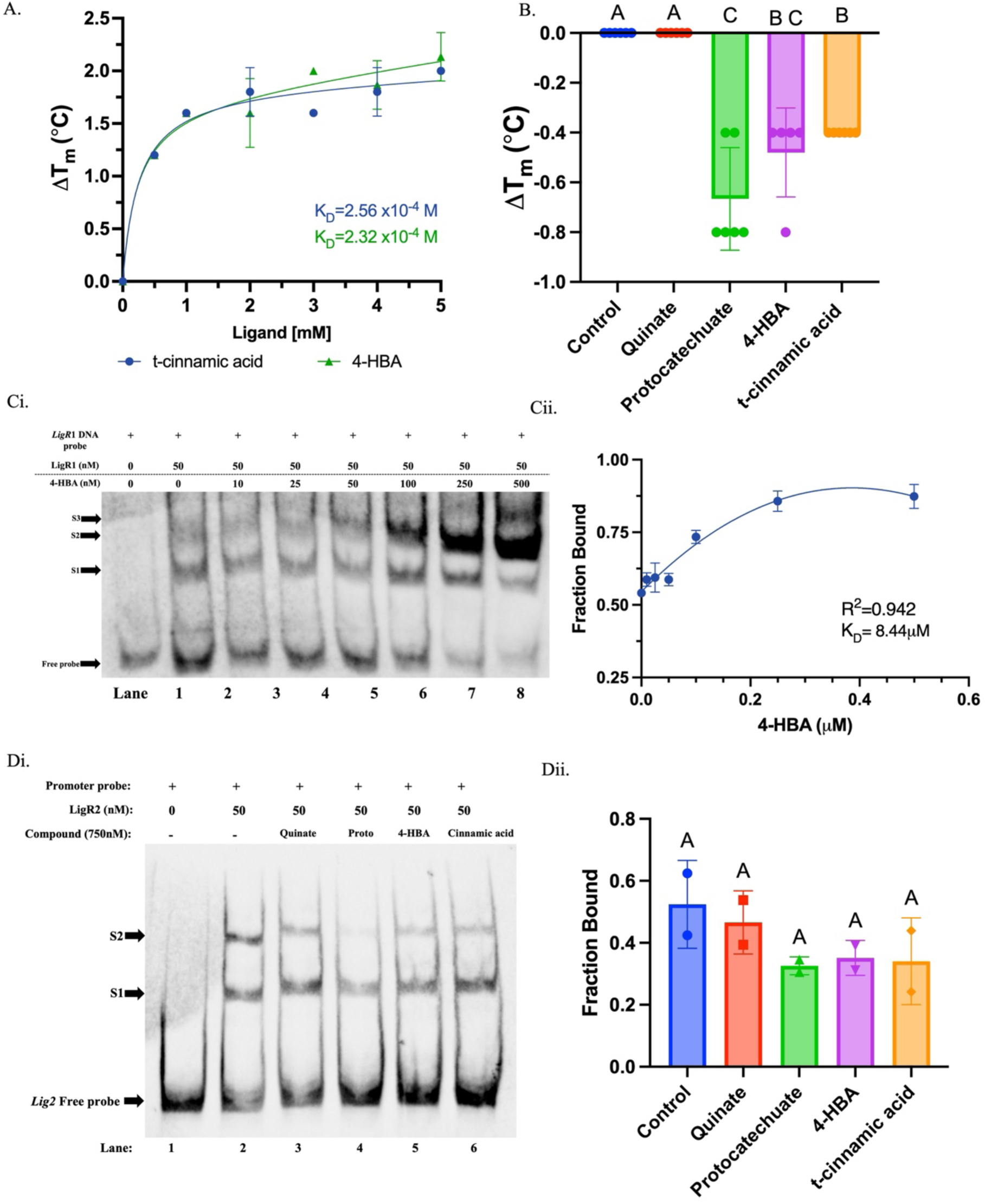
Investigating ligand binding to LigR1 and LigR2 using differential scanning fluorimetry (DSF) and EMSA analyses. A) Saturation curve showing the thermal shift (ΔTm in °C) of LigR1 in response to increasing concentrations (0–5mM) of *t*-cinnamic acid or 4-HBA, with each point representing the average ΔTm per concentration (n = 3–4). B) Thermal shift (ΔTm in °C) of LigR2 in the presence of 0.25mM quinate, protocatechuate, 4-hydroxybenzoate (4-HBA), and *t*-cinnamic acid (n = 5). Ci) EMSA analysis of LigR1 binding to the *lig1* promoter with increasing concentrations of 4-HBA (0–500nM). Lanes 1–7 correspond to 4-HBA concentrations of 0, 10, 25, 50, 100, 250, and 500nM. LigR1 protein and *lig1* promoter probe concentrations were 50nM and 15ng·μL⁻¹, respectively. Mobility shifts are labeled as Shift 1 (S1), Shift 2 (S2), and Shift 3 (S3). Cii) Quantification of LigR1 binding to the *lig1* promoter in response to increasing concentrations of 4-HBA, shown as the fraction of DNA bound based on the ratio of band intensities between bound and unbound DNA. Di) EMSA analysis of LigR2 binding to the *lig2* promoter in the presence of 750nM of each ligand: quinate, protocatechuate (proto), 4-HBA, and *t*-cinnamic acid. Dii) Quantification of *lig2* promoter binding by LigR1 in response to increasing concentrations of 4-HBA, based on band intensity analysis as described above. LigR1 protein and *lig1* promoter probe concentrations were 50nM and 15ng·μL⁻¹, respectively. Shift 1 (S1) and Shift 2 (S2) indicate the observed mobility shift positions in the EMSA gel. Data were fitted to the non-linear equation: Y = Bmax/(K_D_ + X) + NSX, where Bmax is the maximum fraction bound and NS the slope of the non-linear component, representing non-specific binding. The coefficients of determination (R^2^) and K_D_ values obtained are shown in the appropriate graph. EMSAs were performed in duplicate. Statistical analysis was performed using one-way ANOVA with Tukey’s multiple comparisons test. Compact letter displays above each bar represent statistically significant differences at a 95% CI. Error bars indicate the SEM.

Next, we investigated how these aromatic metabolites affect LigR1 and LigR2 DNA-binding using EMSA analysis. In the absence of 4-hydroxybenzoate we observed the S1 and S2 bands for LigR1 (Figure 6Ci, Lane 2). Between 250nM to 500nM 4-hydroxybenzoate, the intensity of the S2 band becomes saturated and the S3 band appeared. This contrasted with previous experiments shown in Figure 1Bi, which required at least 240nM protein before the S3 complex could be visible. This is also depicted in the quantification of the bound DNA fraction, which increased by 38% from 0.54 in the control to 0.87 at 500nM 4-hydroxybenzoate (Figure 6Cii). These observations indicate that 4-hydroxybenzoate enhanced LigR1 binding to the *lig1* promoter. For LigR2, we tested the effect of quinate, protocatechuate, 4-hydroxybenzoate, and *t*-cinnamic binding to the *lig2* biotinylated intergenic region (Figure 6Di). These compounds resulted in a decrease in intensity of the S1 and S2 bands, with an increase of the unbound free probe (Figure 6Dii). Overall, the binding of these compounds to LigR2 cause it to dissociate from the intergenic region as a mechanism to relieve its repressive action as seen from the *β*-galactosidase reporter assay.

### Modelling the ligand binding pocket of LigR1 and LigR2

The crystal structure of LigR1 revealed that each monomer (Chain A-D) contained an acetate molecule within the EBD. The carboxylic acid group of the acetate molecule hydrogen bonds with the backbone amine group of S145 and A144. The CH3-group of acetate points downward towards the loop region of the binding pocket (Figure 7A). Although we could not co-crystallize LigR1 with 4-hydroxybenzoate, we reasoned from our DSF and EMSA data that 4-hydroxybenzoate could be accommodated within this region. We positioned 4-hydroxybenzoate within the binding pocket using LigandFit from the Phenix Suite, such that the COOH-group superimposed the acetate molecule, which was already present (Figure 7B). The aromatic ring of 4-hydroxybenzoate was in proximity to a tyrosine, Y200, facilitating π-stacking interaction with the ring. Chain B of LigR1 was overlayed with the putative functional homolog, PobR, from *Acinetobacter baylyi* ADP1 having two substitutions (ΔL141/L220V) in-complexed with 3-HBA (PDB: 5HPI)^36^. We used this PobR for structural comparison since it has the highest Z-score (26.2) (Figure 7B). The position of the 3-HBA in PobR closely corresponded to the position of acetate in LigR1 and superposed with the modeled 4-hydroxybenzoate. Our structural analysis and DSF results indicated that 4-hydroxybenzoate can serve as an effector molecule for LigR1.

**Figure 7.**
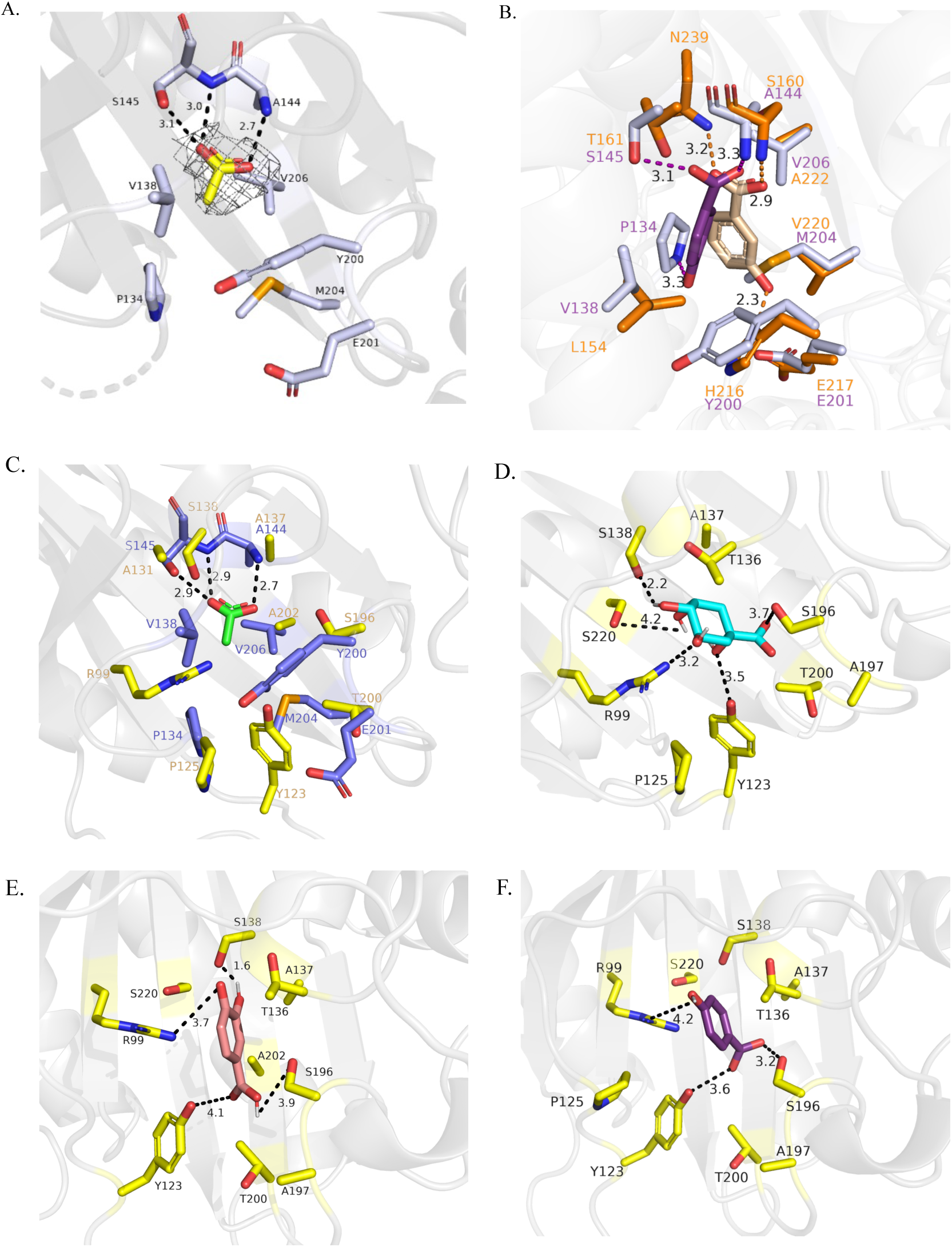
Active site analysis of LigR1 crystal structure and LigR2 AlphaFold Model. A) Binding site residues of LigR1 Chain B with acetate in the ligand binding pocket of the C-terminal EBD. Acetate is depicted in yellow, and the surrounding residues are in light purple. The dashed lines depicted H-bonding interactions and their corresponding length in Å. Mesh around the acetate molecule shows the electron density observed B) Superimposition of LigR1 Chain B with the PobR variant from *A. baylyi* ADPI complexed with 3-HBA (PDB ID: 5HPI). 3-HBA is highlighted in cyan, and its AS residues are depicted in light pink. 4-hydroxybenzoate was manually position within the LigR1 active site to align with the carboxylate group of acetate. Purple residues correspond to the LigR1 structure. One letter residue code shows the positioning of the residue within the sequence. C) Superimposition of LigR1 (purple) and LigR2 (yellow) active site residues in complex with acetate (green). D-F) Modelling quinate, protocatechuate, and 4-hydroxybenzoate ligand binding using AutoDock, respectively. All images were generated from PyMOL version 3.1.6.1

Next, we superimposed the acetate bound LigR1 structure against the LigR2 AlphaFold model to compare the ligand binding residues between the two transcription factors (Figure 7C). Between the two structures, there is strong conservation among the residues coordinating hydrogen bonding with the acetate carboxylate group. Since we identified in our *β*-galactosidase reporter assay that LigR2 operon expression is activated by quinate, protocatechuate, and 4-hydroxybenzoate, we modelled the metabolites within the identified binding site using AutoDock. This analysis showed a calculated affinity value (kcal/mol) of −1.464 for quinate (Figure 7D), −2.623 for protocatechuate (Figure 7E), and −3.233 for 4-hydroxybenzoate (Figure 7F). Electrostatic surface potential maps of LigR1 and LigR2 indicates the primary ligand binding pockets are both enriched in positively charged amino acids (Supplementary Figure 5). However, the size of the binding pockets are vastly different between LigR1 and LigR2 thus providing some mechanistic insight behind LigR2s broad ligand specificity (Supplementary Figure 5C-F).

### LigR1 and LigR2 intricately modulate glucose and aromatic carbon compound utilization

To investigate the biological role of LigR1, we generated an unmarked deletion of the *ligR1* coding region using homologous recombination. For complementation, the full-length *ligR1* gene was expressed under the *lacZa* promoter in the Δ*ligR1* strain, designated Δ*ligR1*:LigR1. We also obtained a transposon mutant of *pp_2605* (Δ*oxred2*), which contains a kanamycin resistance cassette disrupting the gene. We first monitored the growth of WT, Δ*ligR1*, *ΔligR1*:LigR1, and *Δoxred2* strains in M9 minimal media supplemented with 25mM glucose over 24 hours (Figure 8A). The WT strain reached a final OD600nm of 3.52, while Δ*ligR1* exhibited severe growth impairment reaching only 0.197 at OD600nm. The Δ*oxred2* strain showed a milder defect, achieving an OD600nm of 2.0. The *ΔligR*1:LigR1 complemented strain displayed full growth recovery with 0.4mM IPTG, and was used in all subsequent studies involving IPTG. Growth kinetics revealed that WT cells and the complemented strain entered exponential phase after 4 hours. In contrast, *Δoxred2* exhibited a delayed exponential phase onset up to 8 hours, while Δ*ligR1* failed to enter exponential growth entirely. Quantification of the AUC from growth data further supported these observations (Figure 8B). Strikingly, the Δ*ligR1* strain exhibited an unusual aggregation phenotype, forming visible clumps as early as 3–5 hours, which became more pronounced over time. To determine whether this was linked to cell lysis, we measured protein content in the supernatant using Bradford assays (Supplementary Figure 6). The Δ*ligR1* culture had a higher OD595/OD600nm ratio than the WT or complemented strains, indicating increased cell lysis and thus protein content in the culture media.

**Figure 8.**
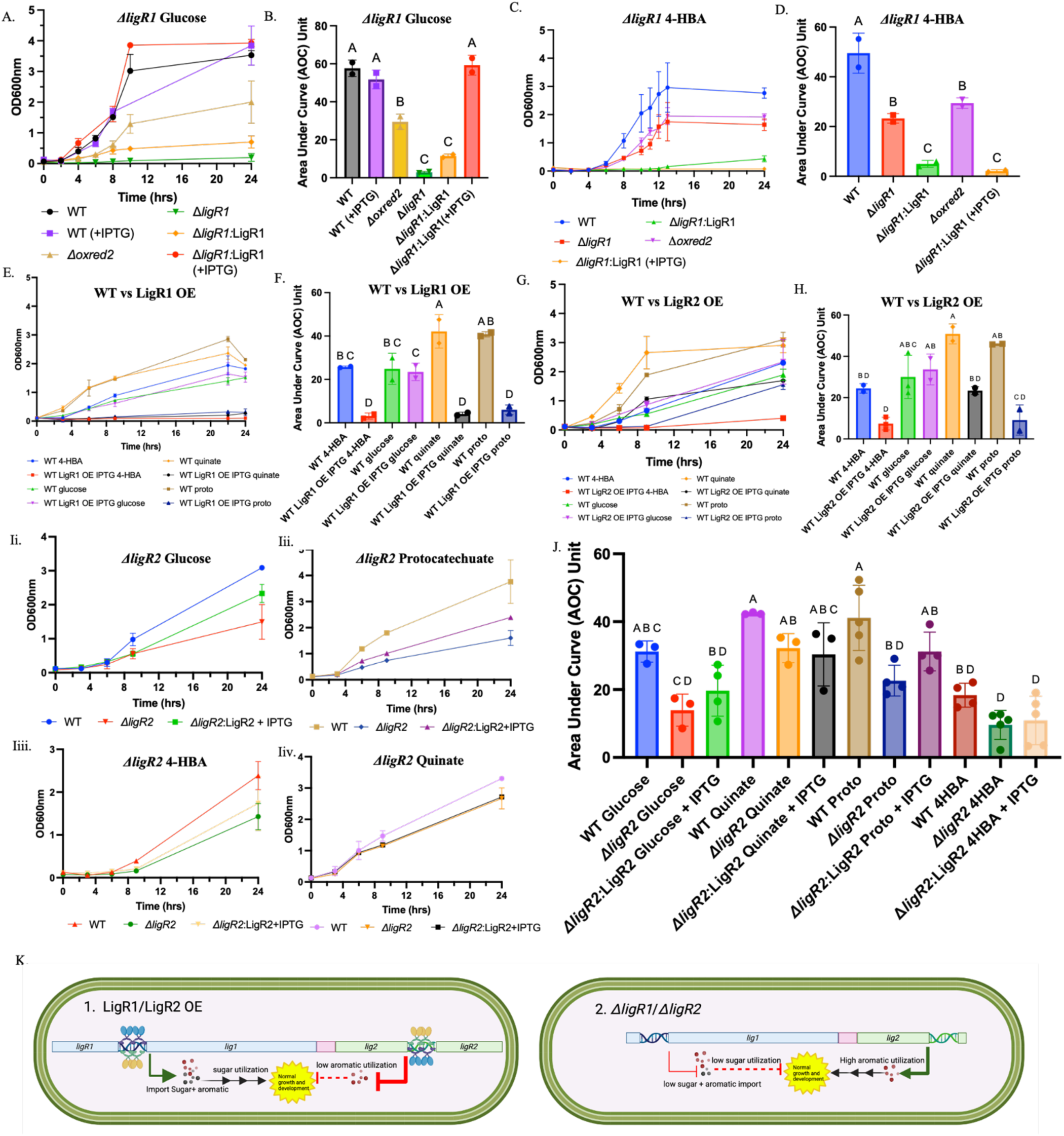
Growth analysis of *P. putida* KT2440 under regulation by LigR1 and LigR2. A) Growth curves of WT (black), WT + IPTG (purple), *Δoxred2* (brown), *ΔligR1* (green), *ΔligR1*:LigR1 complement (orange), and *ΔligR1*:LigR1 + 0.4mM IPTG (red) strains in M9 minimal media supplemented with 25mM glucose, monitored by OD600nm over 24 hours (n= 2). B) Average quantification of AUC for the glucose growth curves shown in panel A. C) Growth curves of WT (blue), *Δoxred2* (purple), *ΔligR1* (red), *ΔligR1*:LigR1 complement (green), and *ΔligR1*:LigR1 + 0.4mM IPTG (orange) in M9 minimal media with 25mM 4-HBA, monitored over 24 hours (n= 2). D) Average AUC quantification for the 4-HBA growth curves shown in panel C. E) Growth curves of WT and LigR1-overexpressing strains (LigR1 OE) in M9 minimal media supplemented with 25mM of either glucose, 4-HBA, protocatechuate (proto), or quinate, with 0.4mM IPTG. F) Average AUC quantification for WT and LigR1 OE growth across the different carbon sources. G) Growth curves of WT and LigR2-overexpressing strains (LigR2 OE) in M9 minimal media with the same four carbon sources as in panel E and IPTG H) Average AUC quantification for WT and LigR2 OE growth curves across all tested carbon sources. I) Growth curves of WT, *ΔligR2*, and *ΔligR2*:LigR2 complemented strains in M9 minimal media with 25mM of glucose (Ii), protocatechuate (Iii), 4-HBA (Iiii), and quinate (Iiv), monitored over 24 hours (n = 3–5). J) AUC quantification for the growth curves shown in panel I across all carbon sources. Statistical analysis was performed using one-way ANOVA followed by Tukey’s multiple comparisons test. Compact letter display denote statistically significant differences at a 95% CI. Error bars represent the SEM. K) Schematic of the growth analysis of *P. putida* KT2440 under regulation by LigR1 and LigR2.In the LigR1 or LigR2 overexpression strain, import of sugars and aromatics is facilitated by expression of the MFS transporter from the *lig1* operon enabling normal growth and development. In contrast, LigR1 and LigR2 overexpression results in attenuation of the *lig2* operon and prevent utilization of the aromatic compounds resulting in growth suppression. In the Δ*ligR1* strain, the *lig1* operon expression is attenuated resulting in low sugar and aromatic import and growth suppression in glucose. In the Δ*ligR2* strain, the *lig2* operon is highly expressed and results in aromatic utilization for normal growth and development.

We next tested growth in M9 minimal media with 25mM 4-hydroxybenzoate (Figure 8C). The WT strain reached an OD600nm of 2.76 after 24 hours, while Δ*ligR1* and Δ*oxred2* achieved OD600nm of 1.6 and 1.9, respectively. The complemented strain, however, showed unexpectedly poor growth, reaching an OD600nm of only 0.075 with IPTG and 0.436 without IPTG. We investigated the effect of LigR1 overexpression by engineering WT cells to also overproduce *ligR1* under IPTG induction (Figure 8E). Remarkably, LigR1 overexpression strongly suppressed *P. putida* growth on the following substrates: quinate, 4-hydroxybenzoate, and protocatechuate. The most severe inhibition occurred with 4-hydroxybenzoate, where the cells failed to grow. Further, we established that the growth rate of the WT and LigR1 overexpression *P. putida* cells are similar over a 24hr period in glucose.

We also examined the effects of LigR2 overexpression and compared its growth phenotype to that of LigR1 (Figure 8G). Unlike the severe suppression seen with LigR1, LigR2 overexpression resulted in milder growth inhibition. On 4-hydroxybenzoate, WT cells reached an OD600nm of 2.3, while LigR2-overexpressing cells grew only to 0.4 OD600nm. Growth on quinate and protocatechuate was less affected: WT cultures achieved OD600nm values of 2.9 and 3.1, respectively, whereas LigR2-overexpressing strains reached OD600nm of 1.69 and 1.55. In contrast, LigR2-overexpressing cells grown in glucose reach an OD600nm of 0.2583 compared to the WT cells which grew to a final OD600nm of 1.89 in 24hrs. Quantification of the area under the curve support our growth curve observations for Δ*ligR1* grown in 4-hydroxybenzoate, LigR1 and LigR2 overexpression (Figure 8D,F,H).

Finally, we investigated the effects of disrupting Δ*ligR2* expression on growth in glucose, quinate, protocatechuate, and 4-hydroxybenzoate (Figure 8I-J). A significant difference was observed between the WT and Δ*ligR2* growth curves only in condition supplemented with protocatechuate. The WT cells reached a final mean OD600nm of 3.76, while the Δ*ligR2* cells reached a mean OD600nm value of 1.6, suggesting that Δ*ligR2 P. putida* growth on protocatechuate is negatively impacted. Overall, this data indicates that manipulating expression of LigR1 and LigR2 transcriptional regulators drastically impacts bacterial growth physiology. In particular, overexpression of LigR1 and LigR2 result in growth suppression on aromatic compounds. While mild phenotypes were observed for aromatic compound utilization by Δ*ligR1* and Δ*ligR2,* glucose utilization by Δ*ligR1* was severely impacted

### *ΔligR1* alters bacterial cell morphology and biofilm formation

Amorphous cell aggregation was observed during growth analysis which led us to investigate cell morphology using confocal microscopy. The *ΔligR1* mutant exhibited distinct morphological changes compared to WT cells in LB media (Figure 9Ai). While WT cells maintained their characteristic short rod shape appearance, *ΔligR1* cells were very elongated. Quantitative analysis using ImageJ revealed the *ΔligR1* cells averaged 3.1μm in length, exactly double that of WT cells (Figure 9Bi). This elongation phenotype was even more pronounced when cells were grown in M9 minimal media supplemented with glucose, where *ΔligR1* cells measured 4.94μm compared to 2.79μm for WT (Figure 9Bii). Complementation of *ΔligR1* with LigR1 restored cell length to near-WT levels (1.5μm), while the *Δoxred2* mutant showed an opposite phenotype, with shorter (1.08μm), more rounded cells. The *ΔligR1* mutant displayed striking cell-cell adhesion, forming complex aggregates where individual clusters are observed to be long chains of bacteria resembling branched networks. This aggregation phenotype correlated with the visible clumping observed in liquid cultures. Neither WT nor the complemented strains showed this behavior (Figure 9Aii). The *Δoxred2* mutant showed no significant cell shape alteration from WT and only minimal evidence of clumping at 4x magnification. When grown in 4-hydroxybenzoate supplemented media, *ΔligR1* cells maintained their distinctive aggregation pattern (Figure 9Aiii). A subpopulation of these cells showed extreme elongation compared to WT, with these morphological anomalies clearly visible in the 4x zoom panels. The complemented strain again showed near-complete restoration of normal morphology under these conditions. The *Δoxred2* mutant showed no significant morphological changes in 4-hydroxybenzoate compared to glucose conditions. This data further supports the specific role of LigR1 in maintaining proper cell structure during growth on carbon sources.

**Figure 9.**
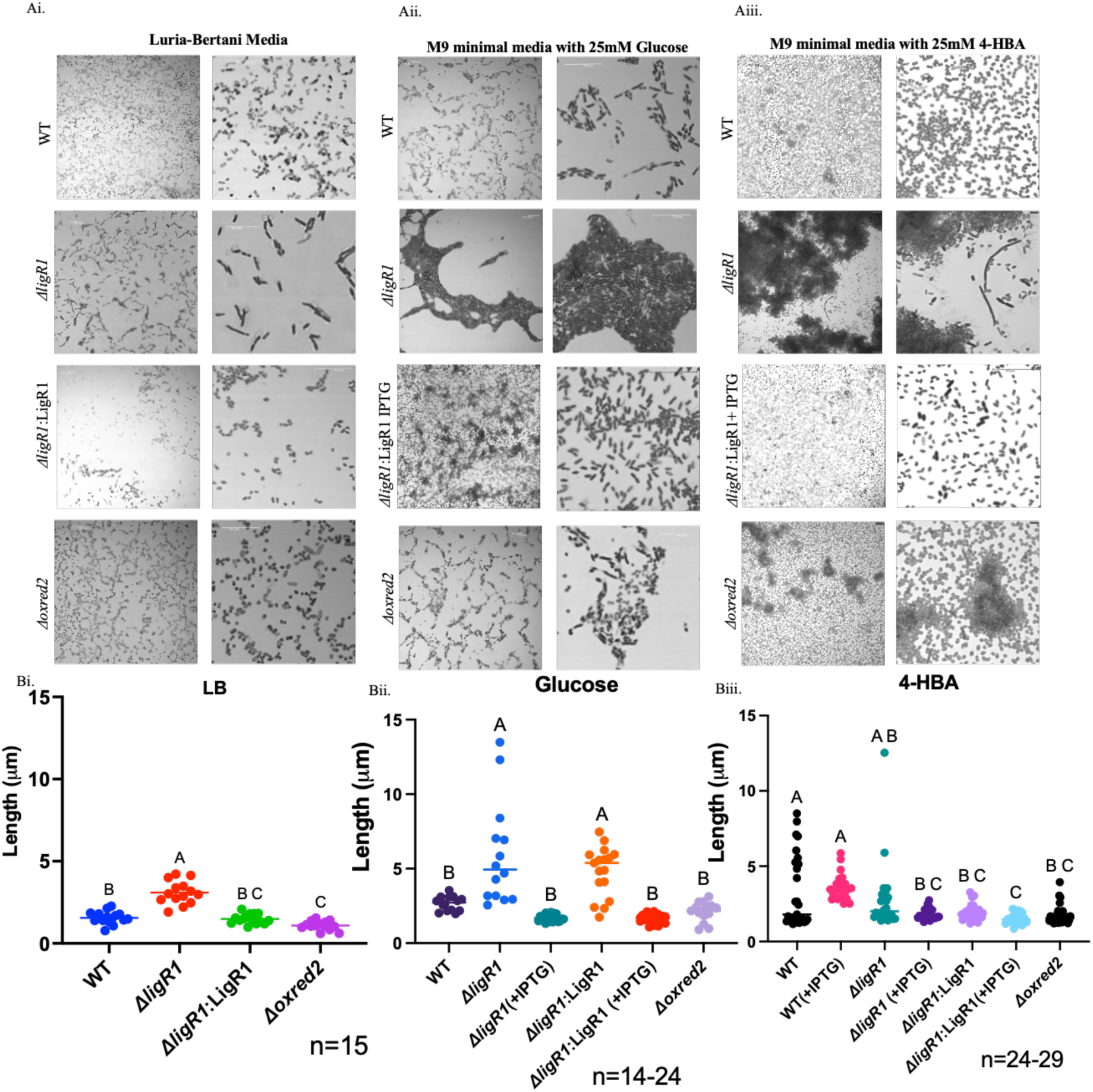
Phenotypic analysis of *P. putida* KT2440 using brightfield confocal microscopy. Ai) WT, *ΔligR1*, *ΔligR1*:LigR1, and *Δoxred2* cells were grown in LB statically for 24hrs in sterile petri dishes at 30°C containing glass coverslips. The glass coverslips were stained with 1% crystal violet and imaged using the brightfield confocal microscope at 100x oil immersion. Images were taken at 1x zoom and 4x zoom in a region of interest. Scale for each image was set to 10μm. Aii) WT, *ΔligR1*, *ΔligR1*:LigR1, and *Δoxred2* cells were grown in M9 minimal media with 25mM glucose in the presence or absence of 0.4mM IPTG, processed, and imaged as described above. Aiii) WT, *ΔligR1*, *ΔligR1*:LigR1, and *Δoxred2* cells were grown in M9 minimal media with 25mM 4-hydroxybenzoate pH7.5 in the presence or absence of 0.4mM IPTG, processed, and imaged as described above. Bi-iii) Quantification of bacterial lengths (μm) cells grown in LB, M9 minimal media with 25mM glucose or 4-hydroxybenzoate from confocal images using Fiji (ImageJ). Statistical analysis was conducted by One-Way ANOVA with Tukey’s multiple comparisons tests, represented by compact letter display at 95% CI. Error bars indicate the SEM. Experiments were conducted in two replicates with each point representing the length of a single bacterial cell. N value indicates the number of bacterial cells counted.

## Discussion

The rhizosphere is a chemically rich soil environment with an abundance of aromatic and hexose sugar molecules, which provide nutrients to microorganisms. Although seemingly inconsequential, this chemical complexity imposes a huge challenge for bacteria to constantly prioritize which compounds to metabolize. This decision directly impacts these organisms ability to continuously compete for dominance in the rhizosphere. Studies have shown that metabolic prioritization is primarily determined by the carbon catabolite repression (CCR) system by inhibiting the translation of mRNA from genes involved in the metabolism or transport of unpreferred carbon sources until preferred ones are depleted^6^. The CCR system is controlled by the Crc with Hfq proteins by forming a ribonucleoprotein complex, which bind to the 5’end of mRNA at an attachment site defined by the sequence AANAANA ^67–69^. This form of global regulation is energetically costly as it relies on continuous mRNA synthesis for proteins all of which are not utilized by the organism at a given time. In addition to CCR, transcription factors are important in delegating how gene expression within operons are modulated in respect to the relative abundance of specific compounds in the environment. These studies, however, offer limited insight into metabolic prioritization, as they focus primarily on a single compound or specific class of compounds processed through one pathway. In this study, we have identified two adjacent, *lig1* and *lig2* operons in *P. putida* KT2440, which serve bifunctional roles to enable selective import and utilization of glucose, shikimate pathway intermediates and aromatic compounds through coordinated regulation by the IclR-family regulators LigR1 and LigR2.

In our quest to understand the interplay between LigR1 and LigR2 we undertook a multidisciplinary approach to investigate their role in *P. putida* KT2440. This approach consisted of gene expression analyses to investigate their regulatory function, as well as biochemical and microbial growth assays to establish the mechanism and biological impact on bacterial physiology. These studies provided valuable insights into the interplay between the two transcription factors in modulating selective import and utilization of related types of polyol chemical compounds.

The regulation of *lig1* and *lig2* operons gene expression in response to different nutrient availability is interconnected with both LigR1 and LigR2 transcriptional regulators. This is consistent with the fact that some downstream chemical compounds produced by proteins from the *lig1* operon are subsequently utilized by proteins from the *lig2* operon. This is supported by data from the gene expression and microbial growth analyses which established that LigR1 is required for *lig1* operon expression, as deletion of LigR1 attenuates *lig1* gene expression and inhibits *P. putida* growth on glucose. Interestingly, LigR1 overexpression has a stronger repressive effect on gene expression from *lig2* in contrast to *lig1* operon gene expression. This repressive effect of LigR1 on *lig2* gene expression was further observed with suppressed growth on certain aromatics but restored growth on glucose in the *P. putida* LigR1 overexpression strain. Unexpectedly, we observed that LigR2 overexpression stimulated *ligR1* expression, but did not affect *lig1* operon expression or growth of the organism on glucose. This regulatory interplay between LigR1 and LigR2 was further observed with the deletion of *lig2* which reduced *lig1* operon expression thus resulting in impaired ability of the organism to grow on glucose. Expression of the *lig2* operon was stimulated by specific aromatics, with induction strongly enhanced by *ligR2* deletion. Despite this, we observed a mild decrease in growth on the aromatics, presumably due to attenuation of the *lig1* operon which contains the MFS transporter. Finally, LigR2 overexpression attenuates *lig2* operon expression, correlating with growth suppression on aromatics. Our EMSA studies further demonstrated LigR1 and LigR2 participate together in the regulation of these operons, as LigR2 also exhibited binding to the *lig1* intergenic region at conserved TAAT/ATTA motifs. LigR1 and LigR2 formed three distinct DNA–protein complexes at their shared intergenic region, for which we speculate they represent dimeric, tetrameric, and higher-order oligomeric states, while only a single complex was observed for LigR2 at the *lig1* intergenic region. The biological relevance of these DNA-protein oligomeric states is not immediately evident and thus will require additional studies. Mutational analysis of the *lig1* promoter revealed that introducing a GCTG motif reduced LigR1 binding but enhanced LigR2 binding affinity, suggesting these regulators use distinct DNA recognition mechanisms. Drawing from structural studies of IclR-family and AraC/XylS-family transcription factors, we propose that LigR1 relies on A/T-rich sequences that promote DNA bending for stable tetramer formation, while LigR2 favors G/C-rich sequences that enhance rigidity and groove widening to enable alternative binding modes^70^.

An understanding of the genetic composition of *lig1* and *lig2* operons and how they relate to *P. putida* dominance in the rhizosphere is of tremendous importance. The presence of the PP_2604 Lig1 MFS transporter in the *lig1* operon suggested it functions upstream of the *lig2* operon to enable the selective import of polyol compounds including hexose sugars such as glucose, shikimate pathway intermediates and aromatic compounds. PP_2604 shows high sequence homology to PcaK, a known protocatechuate/4-hydroxybenzoate transporter. Additional bioinformatic analysis revealed relatedness to putative hexuronate/glucarate transporter such as the *E. coli* D-galactonate:H+ symporter (PDB ID: 6e9n, Z-score: 2.8, RMSD: 400). Although, *P. putida* does not metabolize D-galactonate, structurally similar sugars like glucose or six-membered ring compounds like quinate may be recognized through their conserved −OH groups. This is not surprising since this MFS belongs to the polyol family of transporters which are shown to transport compounds decorated with multiple −OH groups including common sugars and benzoic acid derivatized aromatic compounds.

Within the *lig1* operon, we also identified RifI, a shikimate dehydrogenase homolog closely related to AroE, the well-characterized enzyme of the shikimate pathway involved in aromatic amino acid biosynthesis. While AroE’s function is well established, RifI remains functionally uncharacterized despite extensive structural investigation^42,71^. The presence of RifI in *P. putida* is unusual compared to other bacteria such as *Amycolatopsis mediterranei* and *Streptomyces coelicolor*^72–75^. In these organisms, RifI resides within a genomic operon encoding enzymes of the aminoshikimate pathway, which synthesize 3-amino-5-hydroxybenzoic acid, an essential precursor for ansamycin and mitomycin antibiotic production. Based on *P. putida* RifI gene cluster, we hypothesize that it is involve in diverting shikimate pathway intermediates from chorismate biosynthesis towards protocatechuate for entry into the *β*-ketoadipate pathway or the meta-cleavage pathway. This is consistent with the interconnectivity between *lig1* and *lig2* operons, in which *lig1* is involve in sugars and aromatics import and *lig2* encodes proteins for aromatics metabolism. Further, this hypothesis is supported by studies in *L. monocytogenes* which investigated the role of the *qui1* and *qui2* operons regulated by the LysR-type QuiR/QuiR2^44,46^. The *qui1 and qui2* operons mediate shikimate and quinate uptake and conversion to protocatechuate through sequential oxidation and dehydration reactions^44,46^. We propose *P. putida lig1/lig2* operons employs a similar metabolic framework to *qui1/qui2* for alternative utilization of shikimate pathway intermediates, whereby quinate and shikimate-derived monolignols are diverted away from chorismate biosynthesis and instead used for alternative roles-in this case for energy-in a metabolically rich rhizosphere environment where *P. putida* is frequently associated.

We expected that *ΔligR1* mutant impairment in glucose and aromatic compounds will impact growth ability of the organism however surprisingly, we observed distinct cell aggregation in liquid culture. Analysis of the cell aggregates revealed that these growth defects were accompanied by striking cell elongation and aggregation phenotypes. In light of this observation, we propose that disruption of the MFS transport system through *ΔligR1* mutation leads to impairment of energy supply for optimal production of structural elements of the cell wall and to support cell division. The observed filamentation and aggregation phenotypes parallel observations in *Mycobacterium smegmatis*, where the IclR-family regulator Rv2989 induces similar morphological changes during dormancy^77^. While *P. putida* does not enter dormancy, the metabolic disruption caused by *ΔligR1* appears to trigger these analogous responses when transport is impaired. This elongation phenotype is reminiscent of *P. aeruginosa* under nutrient limitation as an adaptation to increase surface area for nutrient scavenging^78^. Our cell lysis experiments showed increased protein content in the growth media under glucose conditions, suggesting *ΔligR1*-mediated aggregation represents a compensatory mechanism for impaired nutrient uptake including an underdeveloped cell wall. In addition, it is expected that cell lysis of *ΔligR1* mutant contributes extracellular DNA (eDNA) to the establishment of the biofilm matrix. For example, in *P. aeruginosa, streptococci* (*S. intermedius, S. mutans*), and *E. faecalis*, eDNA provides structural support by mediating cell-to-cell adhesion through interactions with surface adhesins like type IV pili, and provides nutrients^79^. During biofilm development release of eDNA occurs by two different bacterial subpopulations with varying lysis susceptibility. Some cells may undergo programmed cell death through autolytic pathways as seen in *S. aureus* LytM, while others may employ fratricidal mechanisms using bacteriocins^80,81^. In our *ΔligR1* mutant, we propose that impaired carbon transport triggers altruistic lysis pathways that release eDNA to stabilize biofilms while recycling cellular components, representing an adaptive strategy to cope with metabolic limitations. Further studies are needed to fully elucidate these mechanisms.

Based on our findings, we put forth a model to describe the bifunctional role of the *lig1* and *lig2* operons for both glucose and aromatic compound utilization, which provides *P. putida* a unique opportunity to adapt to fluctuating chemical compounds within the rhizosphere (Figure 10). Under conditions where the rhizosphere is more abundant in hexose sugars, we envision LigR1 and LigR2 working together to induce expression of the *lig1* operon to facilitate their import, along with other known glucose transporters such as GtsABCD. Concurrently, a lower abundance of aromatics results in constitutive repression of the *lig2* operon. The imported glucose molecules are metabolized through the Enter-Duodoroff and Embden-Meyerhof-Parnas pathway for production of pyruvate (PYR) and phosphoenolpyruvate (PEP). While PYR is converted to oxaloacetate (OAA) for entry into the TCA cycle, PEP and erythrose-4-phosphate (E4P) from the PPP enter the shikimate pathway for chorismate biosynthesis. Any aromatics that are imported using the *Lig1* MFS are also diverted into the shikimate pathway. As sugar molecules become depleted and the ratio between glucose to aromatics shifts to represent higher abundance of aromatic compounds the *lig2* operon becomes expressed by inactivating LigR2 to enable transcription of genes which divert shikimate pathway intermediates into protocatechuate for entry in the meta cleavage pathway and subsequently the TCA cycle. This intricate system of regulation by LigR1 and LigR2 enable *P. putida* to prioritize compounds that are overrepresented within the rhizosphere and provide a competitive edge to the bacteria with other microorganisms. The loss of LigR1 is detrimental for these organisms as the inability to import nutrients leads to severe cell aggregation and lysis.

**Figure 10.**
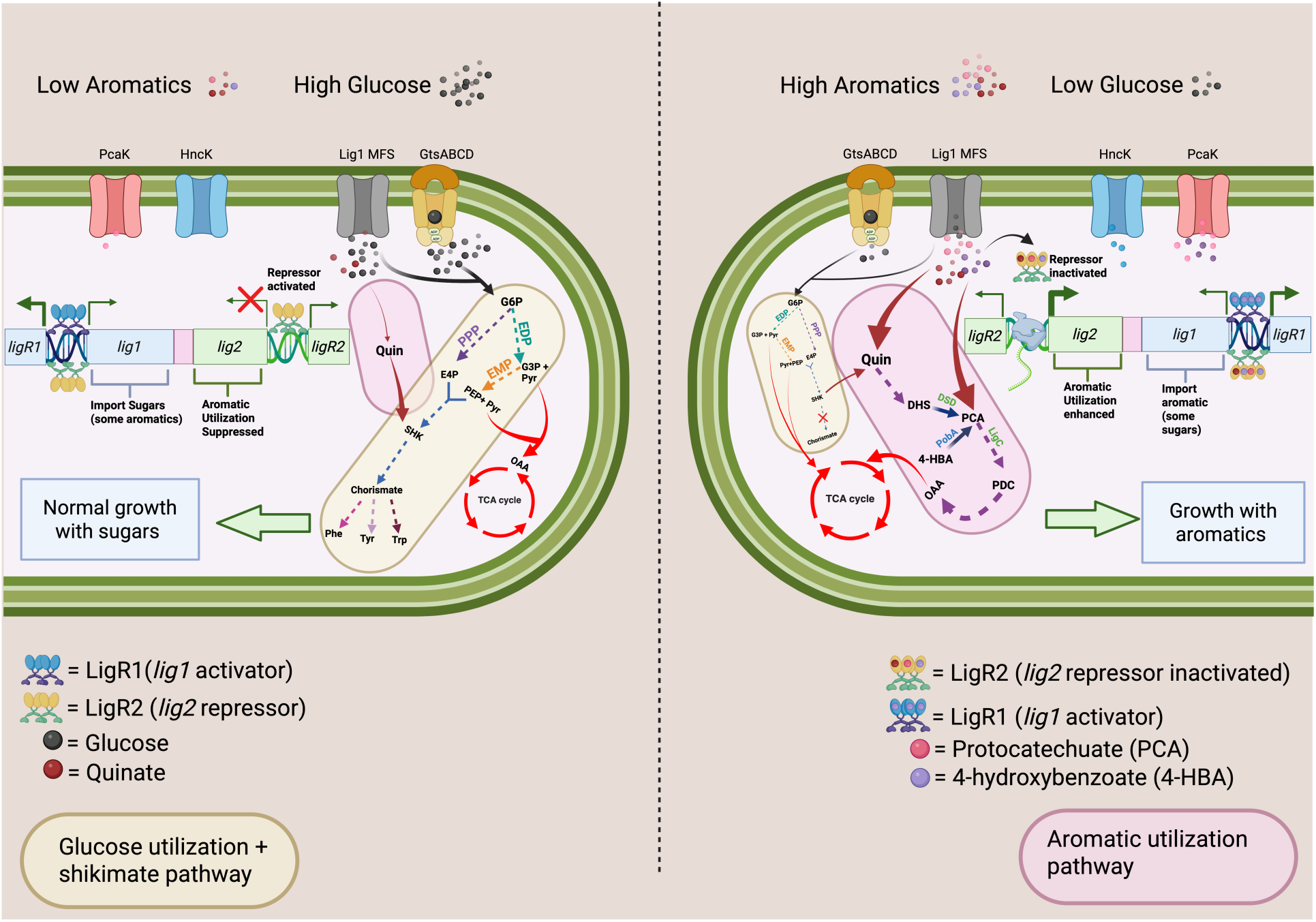
Model of LigR1 and LigR2 mediated regulation in *P. putida* KT2440. Left) Under low aromatics and high glucose, LigR1 and LigR2 activate expression of the *lig1* operon to facilitate import of glucose and some aromatics using the Lig1 MFS transporter (grey). LigR2 also binds to the l*ig2* operon and represses its expression. The GtsABCD transporter together with the Lig1 MFS transporter import glucose, which is converted to glucose-6-phosphate (G6P). G6P enters the Enter-Duodoroff (ED) pathway to yield the end products pyruvate (PYR) and glyceraldehyde-3-phosphare (G3P). G3P also enters the Embden-Meyerhof-Parnas (EMP) pathway which also produces PYR and phosphoenolpyruvate (PEP). The PYR from the ED and EMP pathway is converted to oxaloacetate (OAA) for entry into the tricarboxylic acid (TCA) cycle. G6P also enters the Pentose Phosphate pathway (PPP) which makes erythrose-4-phosphate (E4P). E4P and PEP enter the shikimate pathway for chorismate biosynthesis for downstream production of the three aromatic amino acids: phenylalanine (Phe), tyrosine (Tyr), and tryptophan (Trp). Any import of quinate from the Lig1 MFS is diverted to shikimate, which also feeds into the shikimate pathway. This supports normal growth with sugars. The glucose utilization and shikimate pathway are encompassed by the yellow oval and dominant under high glucose conditions. Right) Under high aromatics and low glucose, the LigR1 and LigR2 regulators continue to express the *lig1* operon to facilitate import of aromatics using the Lig1 MFS transporter. Increasing intracellular concentrations of quinate, protocatechuate, or 4-hydroxybenzoate binds to LigR2 and inactivates repression of the *lig2* operon. The *lig2* operon encoding the putative dehydroshikimate dehydratase (DSD) funnels dehydroshikimate (DHS) into protocatechuate (PCA) intermediate. Subsequently, protocatechuate enters the meta-cleavage pathway where LigC and other enzymes participate in its metabolism to OAA for entry into the TCA cycle. The contributions of glucose for energy and the shikimate pathway are diminished with aromatic utilization pathway dominating for bacterial growth. Protocatechuate and 4-HBA are also imported by the PcaK MFS transporter, located within the pca operon regulated by the IclR-type regulator PcaR. HncK MFS facilitates import of lignin monomers including ferulic acid and p-coumaryl alcohol in *P. putida*, The aromatic utilization pathway encompassed by the red circle dominant under high aromatic conditions, while there is low contribution form the glucose utilization and shikimate pathway.

Overall, our study redefines the metabolic versatility of *P. putida* KT2440 through the discovery of two specialized *lig1/lig2* operons. These operons work together to channel both glucose and shikimate-derived intermediates into energy production via the protocatechuate meta cleavage pathway, broadening the utilization of metabolites within the chemically rich rhizosphere environment. Further, these findings challenge the central dogma of the shikimate pathway as being important for producing the precursor, chorismate, for aromatic compounds biosynthesis to emerging roles in both biosynthesis and for the utilization of carbon compounds for energy through the *β*-ketoadipate pathway. Accumulation of aromatics is sufficient to alleviate repression by LigR2. This metabolic plasticity provides a competitive advantage in the rhizosphere, where the ability to readily adapt to changing metabolic landscapes directly impact the dominance of species. Future studies should explore whether this paradigm extends to other rhizobacteria, and how such pathways might be engineered for biotechnological applications where we can selectively modulate the types of compounds bacteria can metabolize by manipulating the transcriptional machinery that drives the activation or repression of these metabolic genes. Additionally, we provide an alternative mechanism for transcriptional regulation of metabolic prioritization. This mechanism ensures that transcription of the genes within an operon is solely activated with elevated concentration of specific carbon compound.

## Acknowledgements

SMP and DC crystallized LigR1. SP, EK, and DC solved the LigR1 crystal structure. EK designed and conducted the EMSAs, phylogenetics, *β*-galactosidase reporter assays, active site modelling, DSFs, growth curves, confocal microscopy, and BioRender models. SQ conducted EMSAs in Figure 2Ai and Supplementary Figure 2 B,C). DC participated in all aspects in the design of the experiments. EK and DC jointly wrote and edited the manuscript. All BioRender figures were created using the University of Toronto Department of Cell and Systems Biology’s Plan. The presented work was supported by the Natural Science and Engineering Research Council of Canada Discovery Grant (NSERC-DG RGPIN-2020-06052).

