## Supplementary Figures 1-6, Supplementary Table 1-4 for "The interplay between glucose and aromatic compound regulation by two IclR-type transcription factors, LigR1 and LigR2, in *Pseudomonas putida* KT2440"

**Supplemental Figures and Tables:**

| Bacterial Strains | Relevant Characteristics | Purpose | Origin |
| --- | --- | --- | --- |
| ***Escherichia coli*** | | | |
| DH5𝛼 | Plasmid host | | Lab stock |
| DH5𝛼:pET28MOD:*ligR1* | *P. putida* LigR1, Km^R^ | Used to propagate pET28MOD with *P. putida* LigR1 | This study |
| DH5𝛼:pET28MOD:*ligR2* | *P. putida* LigR2, Km^R^ | Used to propagate pET28MOD with *P. putida* LigR2 | This study |
| BL21-CodonPlus (DE3)-RIPL strain | Protein production strain | | Lab Stock |
| BL21:pET28MOD: *ligR1* | *P. putida* LigR1, Km^R^ | Used to express *P. putida* full length LigR1 for protein work | This study |
| BL21:pET28MOD: *ligR2* | *P. putida* LigR2, Km^R^ | Used to express *P. putida* full length LigR2 for protein work | This study |
| ***Pseudomonas putida* KT2440** | | | |
| *P. putida* KT2440 | Wildtype | | Lab stock |
| *ΔligR1* | *ligR1* gene deletion (*pp_2609*) | | This study |
| *ΔligR1*:LigR1 | *P. putida ΔligR1* pBBR1-MCS2-NdeI-l*igR1*, Km^R^ | *ΔligR1* complement | This study |
| *Δoxred2* | *Pp_2605* gene deletion, Km^R^ | | Juan Ramos |
| *ΔligR2* | *ligR2* gene deletion, Chl^R^ ,Km^R^ | | Juan Ramos |
| *ΔligR2:* LigR2 | *P. putida ΔligR2* pucp20g-NdeI-l*igR2*, Chl^R^ , Km^R^, Gm^R^ | *ΔligR2* complement | This study |
| WT LigR1OE | WT *P. putida* pBBR1-MCS2-NdeI-l*igR1*, Km^R^ | Constitutive expression of *ligR1* in WT background | This study |
| WT LigR2OE | WT *P. putida* pBBR1-MCS2-NdeI-l*igR2*, Km^R^ | Constitutive expression of *ligR2* in WT background | This study |
| WT Lig1Op | WT *P. putida* pmp220-*lig1* intergenic region (+23bp *ligR1* and +72bp *rifI)*, Tc^R^ | *lig1* operon expression for 𝛽-galactosidase reporter assay | This study |
| WT Lig1Reg |  | *ligR1* regulator expression for 𝛽-galactosidase reporter assay | This study |
| WT Lig1Op-OE1 | WT *P. putida* pmp220-*lig1* intergenic region (+23bp *ligR1* and +72bp *rifI)*, pBBR1-MCS2-NdeI-LigR1, Tc^R^, Km^R^ | *lig1* operon expression for 𝛽-galactosidase reporter assay with LigR1 constitutively expressed | This study |
| WT Lig1Op-OE2 | WT *P. putida* pmp220-*lig1* intergenic region (+23bp *ligR1* and +72bp *rifI)*, pBBR1-MCS2-NdeI-LigR2, Tc^R^, Km^R^ | *lig1* operon expression for 𝛽-galactosidase reporter assay with LigR2 constitutively expressed | This study |
| *∆ligR1*-1Op | *ΔligR1 P. putida* pmp220-*lig1* intergenic region (+23bp *ligR1* and +72bp *rifI)*, Tc^R^ | *lig1* operon expression for 𝛽-galactosidase reporter assay in *ΔligR1* background | This study |
| *∆ligR1*-1Reg |  | *ligR1* regulator expression for 𝛽-galactosidase reporter assay in *ΔligR1* background | This study |
| Lig2Op | WT *P. putida* pmp220- *lig2* intergenic region (+48bp *ligR2* and +18bp *ligC*) , Tc^R^ | *lig2* operon expression for 𝛽-galactosidase reporter assay | This study |
| Lig2Reg |  | *ligR2* regulator expression for 𝛽-galactosidase reporter assay | This study |
| Lig2Op-OE1 | WT *P. putida* pmp220- *lig2* intergenic region (+48bp *ligR2* and +18bp *ligC*), pBBR1-MCS2-NdeI-LigR1, Tc^R^, Km^R^ | *Lig2* operon expression for 𝛽-galactosidase reporter assay with LigR1 constitutively expressed | This study |
| Lig2Op-OE2 | WT *P. putida* pmp220- *lig2* intergenic region (+48bp *ligR2* and +18bp *ligC*), pBBR1-MCS2-NdeI-LigR2, Tc^R^, Km^R^ | *Lig2* operon expression for 𝛽-galactosidase reporter assay with LigR2 constitutively expressed | This study |
| Lig2Op-EOE | WT *P. putida* pmp220- *lig2* intergenic region (+48bp *ligR2* and +18bp *ligC*), pBBR1-MCS2-NdeI, Tc^R^, Km^R^ | *Lig2* operon expression for 𝛽-galactosidase reporter assay with empty overexpression vector | This study |
| *∆ligR2*-2Op | *ΔligR2 P. putida* pmp220- *lig2* intergenic region (+48bp *ligR2* and +18bp *ligC*) , Chl^R^, Km^R^,Tc^R^ | *Lig2* operon expression for 𝛽-galactosidase reporter assay in *ΔligR2* background | This study |
| *∆ligR2*-2Reg | *ΔligR2 P. putida* pmp220- *lig2* intergenic region (+48bp *ligR2* and +18bp *ligC*) , Chl^R^, Km^R^,Tc^R^ | *LigR2* regulator expression for 𝛽-galactosidase reporter assay in *ΔligR2* background | This study |
| *∆ligR2*-1Op | *ΔligR2 P. putida* pmp220-*lig1* intergenic region (+23bp *ligR1* and +72bp *rifI)*, Chl^R^, Km^R^ ,Tc^R^ | *Lig1* operon expression for 𝛽-galactosidase reporter assay in *ΔligR2* background | This study |
| *∆ligR2*-1Reg |  | *LigR1* operon expression for 𝛽-galactosidase reporter assay in *ΔligR2* background | This study |
| *∆ligR2:*LigR2-2Op | *ΔligR2 P. putida* pmp220- *lig2* intergenic region (+48bp *ligR2* and +18bp *ligC*) , pucp20g-NdeI-LigR2, Tc^R^, Km^R^ , Chl^R^, Gm^R^ | *Lig2* operon expression for 𝛽-galactosidase reporter assay in *ΔligR2:*LigR2 complemented background | This study |
| *∆ligR2:*LigR2-2Reg |  | *LigR2* operon expression for 𝛽-galactosidase reporter assay in *ΔligR2:*LigR2 complemented background | This study |
| *∆ligR2:*LigR2-1Reg | *ΔligR2 P. putida* pmp220-*lig1* intergenic region (+23bp *ligR1* and +72bp *rifI)*, pucp20g-NdeI-LigR2, Tc^R^, Km^R^ , Chl^R^, Gm^R^ | *LigR1* operon expression for 𝛽-galactosidase reporter assay in *ΔligR2:*LigR2 complemented background | This study |
| Lig1-intra | WT *P. putida* pmp220- *Lig1* and *lig2-*150bp intragenic region (+14bp *pp_2604* and +19bp *pp_2603*), Tc^R^ | *Lig1* intragenic operon expression for 𝛽-galactosidase reporter assay between both operons | This study |
| Lig2-intra | WT *P. putida* pmp220- *Lig1* and *lig2-*150bp intragenic region (+14bp *pp_2604* and +19bp *pp_2603*), Tc^R^ | *Lig2* intragenic operon expression for 𝛽-galactosidase reporter assay between both operons | This study |
| **Plasmids** |  |  |  |
| pET28MOD:*ligR1* | pET28MOD *pp_2609-*His6, Km^R^ | Full length LigR1 | This study |
| pET28MOD:*ligR2* | pET28MOD *pp_2601-*His6, Km^R^ | Full length LigR2 | This study |
| pEX18Ap | oriT+ sacB+ gene replacement vector with multiple-cloning site from pUC18; Amp^R^ | Suicide vector for markerless knockout | J.J Harrison |
| pEX18Ap:SOEAD_*ligR1* | oriT+ sacB+ gene replacement vector with multiple-cloning site from pUC18; 300 bp upstream and downstream of *ligR1* Amp^R^ | Suicide vector with homology arms containing 300bp upstream and downstream of *ligR1* for knockout generation | This study |
| pPS858 | Source of the Gm^R^ and FRT cassette Amp^R^, Gm^R^, | Amplify Gm cassette with FRT sites | J.J Harrison |
| pBBFLP | Source of inducible FLP recombinase , Tc^R^ | Excise Gm cassette at FRT sites | J.J Harrison |
| pBBR1-MCS2 | pBBR1 replicon, mob^+^, Km^R^ | Broad-host-range plasmid backbone with multiple cloning site | [Kovach et al., 1995](https://www.sciencedirect.com/science/article/pii/S0167701212002163#bb0055) |
| pBBR1-MCS2-NdeI | pBBR1 replicon, mob^+^, with NdeI cut site in MCS, Km^R^ | Broad-host-range plasmid backbone engineered with an NdeI site at the *lacZ* N-terminus at the multiple cloning site | This study |
| pBBR1-MCS2-NdeI:*ligR1* | pBBR1-MCS2-NdeI:*ligR1* with stop codon, Km^R^ | Broad-host-range plasmid with LigR1 coding sequence under constitutive expression by *lacZ* promoter | This study |
| pBBR1-MCS2-NdeI:*ligR2* | pBBR1-MCS2-NdeI:*ligR2* with stop codon, Km^R^ | Broad-host-range plasmid with LigR2 coding sequence under constitutive expression by *lacZ* promoter | This study |
| pUCp20g | pUCp20 derivative containing constitutive promoter pLac with *Sma*I-flanked Gm^R^ cassette inserted into the unique *Sca*I site within *bla* | Gentamicin-resistant variant of the broad-host range expression vector pUCp20 | West et al., 1994  Centre for the Analysis of Genome Evolution and Function, Toronto, Canada |
| pUCp20g-NdeI | pUCp20g with NdeI cute site in MCS Gm^R^ | Broad-host-range plasmid backbone engineered with an NdeI site at the *lacZ* N-terminus at the multiple cloning site | This study |
| pUCp20g -NdeI:*lig^R^2* | pUCp20g-NdeI:*ligR2* with stop codon, Gm^R^ | Broad-host-range plasmid with LigR2 coding sequence under constitutive expression by *lacZ* promoter | This study |
| Pmp220 | Promoter probe vector, IncQ; Tc^R^ | 𝛽-galactosidase reporter assay | Vittorio Venturi |
| pmp220-lig1op | pmp220-*ligR1* intergenic region (+23bp *ligR1* and +72bp *rifI)*, Tc^R^ | *lig1* operon expression for 𝛽-galactosidase reporter assay | This study |
| pmp220-lig1reg | pmp220-*ligR1* intergenic region (+23bp *ligR1* and +72bp *rifI)*, Tet^r^  pmp220- *lig2* intergenic region (+48bp *ligR2* and +18bp *ligC*) , Tc^R^ | *ligR1* regulator expression for 𝛽-galactosidase reporter assay | This study |
| pmp220-lig2op |  | *lig2* operon expression for 𝛽-galactosidase reporter assay | This study |
| pmp220-lig2reg | pmp220- *lig2* intergenic region (+48bp *ligR2* and +18bp *ligC*) , Tc^R^ | *ligR2* regulator expression for 𝛽-galactosidase reporter assay | This study |
| Pmp220-lig1-intra | pmp220- *Lig1* and *lig2-*150bp intragenic region (+14bp *pp_2604* and +19bp *pp_2603*), Tc^R^ | *Lig1* intragenic operon expression for 𝛽-galactosidase reporter assay between both operons | This study |
| Pmp220-lig2-intra |  | *Lig2* intragenic operon expression for 𝛽-galactosidase reporter assay between both operons | This study |

**Supplementary Table 1. List of bacterial strains and plasmids.**

**Primer Sequences**

| Primer Name | Primer Sequence (5’-3’) | 5’ Modification | Target Gene | Used For |
| --- | --- | --- | --- | --- |
| LigR1-F | GACCATATGAACACACCTGCCATCC | NdeI | *Pp_2609* (no stop codon) | Protein Expression |
| LigR1-R | GTCGGATCCATCCAGCACTGGCCGAAGCA | BamHI |  |  |
| LigR2-F | GCACATATGGCTGGTAGTCAGATCGAACG | NdeI | *Pp_2601* (no stop codon) |  |
| LigR2-R | GTCGGATCCGGCAAACAGCTCGCTGGC | BamHI |  |  |
| LigR1_IR-F | AGGCAGAGGGGCGTGG | Biotin | *lig1*  intergenic region | EMSA |
| LigR1_IR-R | GCATTCGACTCGCTACCAGAGG |  |  |  |
| LigR2_IR-F | GCAGAAGAGCTGTCGGCG |  | *lig2*  intergenic region |  |
| LigR2_IR-R | GGTAATGCTGCCGGCCCA |  |  |  |
| Lig1Op-F | CCTGG GAATTC GGATGGATGGCAGGTGTGTTC | EcoRI | *Lig1* intergenic region (+23bp *ligR1* and +72bp *rifI*) | 𝛽-galactosidase reporter -Operon expression |
| Lig1Op-R | GCCTCTAGAGTTTTGCGGGGACTTCACCTGG | XbaI |  |  |
| Lig1Reg-F | CCTGG GAATTC GTTTTGCGGGGACTTCACCT | EcoRI |  | 𝛽-galactosidase reporter -autoregulation |
| Lig1Reg-R | GCCTCTAGAGGATGGATGGCAGGTGTGTTCAT | XbaI |  |  |
| Lig2Op-F | CCTGGGAATTCGCAGAAGAGCTGTCGGCG | EcoR1 | *Lig2* intergenic region (+48bp *ligR2* and +18bp *ligC*) | 𝛽-galactosidase reporter-Operon expression |
| Lig2Op-R | GCCTCTAGAGGTAATGCTGCCGGCCCA | XbaI |  |  |
| Lig2Reg-F | CCTGGGAATTCGGTAATGCTGCCGGCCCA | EcoR1 |  | 𝛽-galactosidase reporter-autoregulation |
| Lig2Reg-R | GCCTCTAGAGCAGAAGAGCTGTCGGCGG | XbaI |  |  |
| Lig1_2_intra-F | CCTGGGAATTCTATTGGCCGCGGTCTGA | EcoR1 | *Lig1* and *lig2-*150bp intragenic region (+14bp *pp_2604* and +19bp *pp_2603*) | 𝛽-galactosidase reporter expression- *lig2* operon direction |
| Lig1_2_intra-R | GCCTCTAGACAAGGTCGAGAGCGCC | XbaI |  |  |
| Lig2_1_intra-F | CCTGGGAATTCCAAGGTCGAGAGCGCCT | EcoR1 |  | 𝛽-galactosidase reporter expression- *lig1* operon direction |
| Lig2_1_intra-R | GCCTCTAGATATTGGCCGCGGTCTGA | XbaI |  |  |
| FRT-Gm-F | CGAATTAGCTTCAAAAGCGCTCTGA | N/A | Gentamicin | KO generation |
| FRT-Gm-R | CGAATTGGGGATCTTGAAGTTCCT | N/A |  |  |
| LigR_SOEA-F | CGACCGAATTCCAGCGCCGCCTCGTGCAGGTC | EcoRI | +300bp upstream of *ligR1* (+156bp intergenic region +144bp into *rifI*) |  |
| GM-FRT-LigR_SOEB-R | TCAGAGCGCTTTTGAAGCTAATTCGGCATTCGACTCGCTACCAGA | N/A |  |  |
| Gm-FRT-LigR_SOEC-F | AGGAACTTCAAGATCCCCAATTCGCGCAGACATGGCGCAGTTTGC | N/A |  |  |
| LigR_SOED-R | CGAACGGGATCCCGTGTGGGCGATCGCATC | BamHI | +300bp downstream of *ligR1* (+149bp and +151bp into *pp_2610*) |  |
| LigR_KO-F | TGCAACCTACTTGCCGCGC | N/A | +106bp upstream of *ligR1* | KO check |
| LigR_KO-R | TTCAACGTCCGAGCAGCGC | N/A | +404bp downstream of *ligR1* |  |
| pBBR1-MCS2-NdeI-SDM-F | AACAATTTCACACAGGAAACACATATGACCATGATTACGCCA | N/A | pBBR1-MCS2 | Introduce NdeI cut site in MCS at lacZ N-terminus |
| pBBR1-MCS2-NdeI-SDM-R | TGGCGTAATCATGGTCAT ATGTGTTTCCTGTGTGAAATTGTT | N/A |  |  |
| LigR1_Comp-F | GCACATATGAACACACCTGCCATCCATCCC | NdeI | *Pp_2609-*stop | Complementation of *ΔligR1* and overexpression in WT background. |
| LigR1_Comp-R | GTCGGATCCTCAATCCAGCACTGGCCGAAG | BamHI |  |  |
| LigR2_OE-F | GCACATATGGCTGGTAGTCAGATCGAACG | NdeI | *Pp_2601*-stop | LigR2 overexpression in WT background and complementation of *ΔligR2* |
| LigR2_OE-R | GTCGGATCCTCAGGCAAACAGCTCGCTGGC | BamHI |  |  |

**Supplementary Table 2. List of primer sequences.**

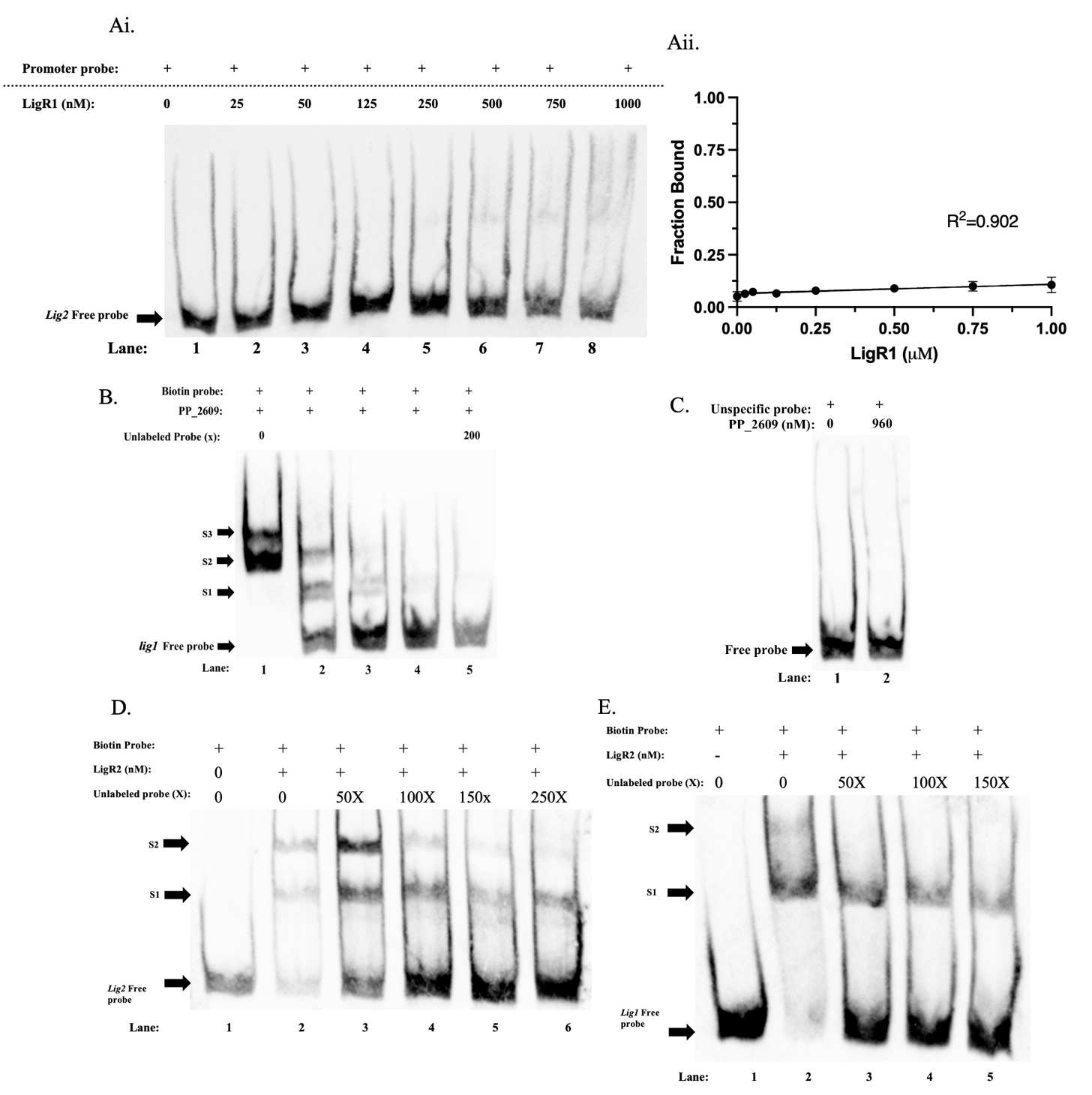

**Supplementary Figure 1.** **Competition** **EMSA’s for LigR1 and LigR2 DNA binding.** Ai) EMSA analysis of LigR1 binding to the *lig2* promoter probe. Lanes 1-8 depict increasing concentrations of LigR1 from 0nM, 25nM, 50nM, 125nM, 250nM, 500nM, and 1000nM. **Aii)** Quantification of *lig2* promoter binding by LigR1, shown as the fraction of DNA bound at each concentration, based on band intensity comparisons between the bound and unbound DNA. B) 256nM of LigR1 incubated with 0-200x unlabeled *lig1* DNA probe for 30 minutes followed by 15ng μL^-1^ of biotinylated *lig1* DNA probe. C) EMSA of LigR1 with a *P. putida* biotinylated glycerol-3-phosphate acetyltransferase gene fragment tested at 0 and 960nM of protein. D) 125nM of LigR2 incubated with 0-250x unlabeled *lig2* DNA probe for 30 minutes followed by 15ng μL^-1^ of biotinylated *lig2* DNA probe. E) 250nM of LigR2 incubated with 0-150x unlabeled *lig1* DNA probe for 30 minutes followed by 10ng μL^-1^ of biotinylated *lig1* DNA probe. Mobility shift positions are labeled as Shift 1 (S1), Shift 2 (S2), and Shift 3 (S3).

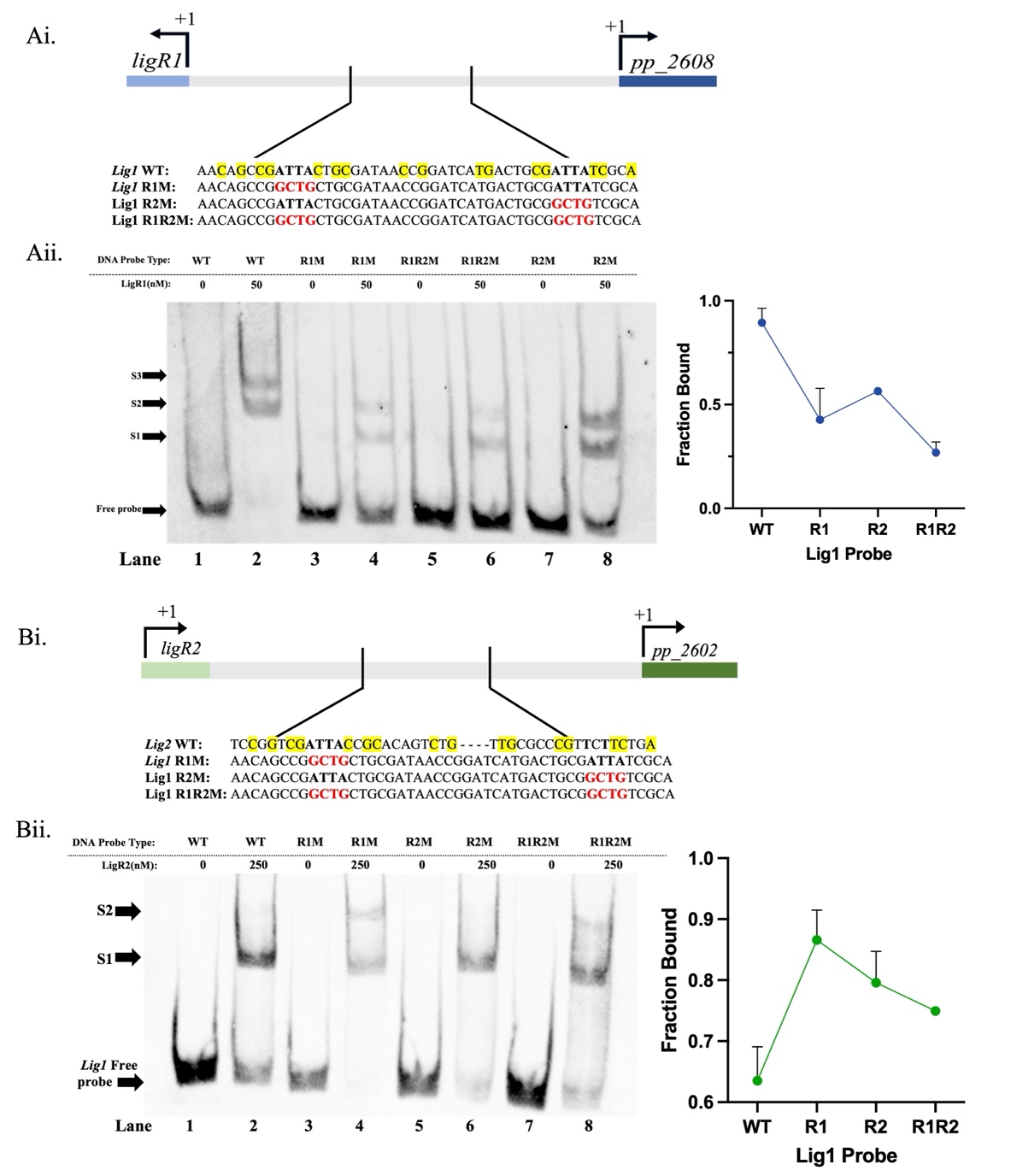

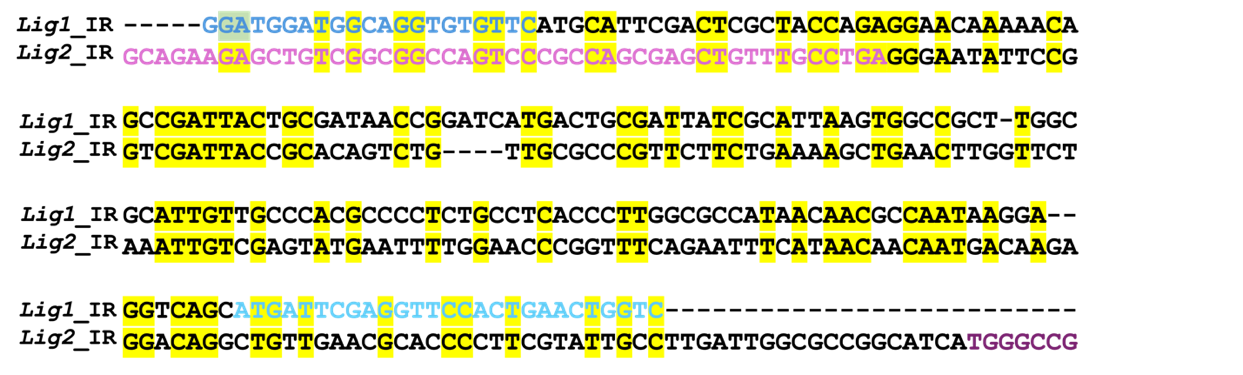

**Supplementary Figure 2. Identification of DNA-binding motifs in the lig1 and lig2 promoter regions.** Ai) Cartoon depiction of the WT lig1 promoter region and synthesized mutants including Region 1 (R1M), Region 2 (R2M), and the combined Region 1 and 2 mutants (R1R2M). WT sequences are indicated with bolded black ‘ATTA’ nucleotides, and mutated sequences are shown as bolded lowercase red ‘gctg’ nucleotides. Yellow highlights denote conserved nucleotides aligning with the Group 2 GTNCG-N₅–₆-CGNAC motif. Each probe is 251 bp long, spanning +23 bp from the ligR1 translation start site and +72 bp from the pp_2608 translation start site. Aii) EMSA of LigR1 binding to 15 ng μL⁻¹ of the biotinylated WT, R1M, R2M, and R1R2M lig1 promoter probes. Lanes 1, 3, 5, and 7 show the migration of the free WT, R1M, R2M, and R1R2M probes without LigR1 protein. Lanes 2, 4, 6, and 8 show the same probes incubated with 50nM LigR1 protein. Shift 1 (S1), Shift 2 (S2), and Shift 3 (S3) indicate the positions of mobility shifts observed. Quantification of the fraction bound for each probe was calculated by comparing band intensities between bound and unbound DNA. Bi) Cartoon depiction of the WT lig2 promoter aligned with the lig1 R1M, R2M, and R1R2M probes, with bolded black ‘ATTA’ indicating WT sequence and bolded lowercase red ‘gctg’ representing the mutated bases. Yellow highlights denote conservation with the Group 2 GTNCG-N₅–₆-CGNAC motif. Bii) EMSA of LigR2 binding to 10 ng μL⁻¹ WT, R1M, R2M, and R1R2M lig1 probes. Lanes 1, 3, 5, and 7 show free probe migration, and lanes 2, 4, 6, and 8 show binding with 50nM LigR2 protein. Mobility shifts S1 and S2 are annotated. Fraction bound was quantified, and binding curves were fitted to the non-linear equation Y = Bmax/(KD + X) + NSX, where Bmax is the maximum fraction bound, NS the slope representing non-specific binding, and KD the dissociation constant. R² and K_D_ values are shown in the plots. EMSA assays were performed in duplicate, and error bars represent the SEM. C) Full nucleotide sequence alignment of the lig1 and lig2 intergenic regions generated using Clustal Omega. Yellow columns represent conserved nucleotides; dashes ‘–’ indicate gaps. Light blue marks the beginning of the rifI sequence, dark blue the start of ligR1, purple the ligC sequence, and pink the ligR2 sequence

| Crystal Parameters | LigR1 |
| --- | --- |
| Space Group | C 1 2 1 |
| Unit Cell (Å^3^) | 178.598 x 69.44 x 95.684 |
| ⍺, 𝛽, 𝛾 (°) | 90, 101.988, 90 |
| Data Collection Statistics |  |
| Wavelength (Å) | 1 |
| Resolution range | 38.09 - 2.11 (2.185 - 2.11) |
| Total reflections |  |
| Unique reflections | 65894 (6541) |
| Completeness (%) | 99.58 (99.89) |
| Mean I/sigma(I) |  |
| Wilson B-factor | 32.04 |
| R_merge_ | 0.11 |
| R_meas_ | 0.132 |
| R_pim_ | 0.071 |
| CC1/2 | 0.986 |
| Refinement and model statistics |  |
| Reflections used in refinement | 65885 (6541) |
| Reflections used for R-free | 3273 (313) |
| R-work | 0.1971 (0.2495) |
| R-free | 0.2530 (0.3370) |
| No. of non-hydrogen atoms | 8424 |
| No. of macromolecules | 7854 |
| No. of ligands | 32 |
| No. of solvent | 538 |
| Protein residues | 993 |
| RMS (bonds) (Å) | 0.008 |
| RMS (angles) (°) | 0.91 |
| Ramachandran favoured (%) | 98.68 |
| Ramachandran allowed (%) | 1.12 |
| Ramachandran outliers (%) | 0.20 |
| Rotamer outliers (%) | 1.08 |
| Clashscore | 5.66 |
| Average B-factor (Å^2^) | 37.45 |
| macromolecules | 37.19 |
| ligands | 41.35 |
| solvent | 40.96 |
| PDB code | **9E6A** |

**Supplementary Table 3. X-ray diffraction data collection and refinement statistics for LigR1.** Statistics for the highest-resolution shell are shown in parentheses.

| Species Name | Accession Number | Annotation |
| --- | --- | --- |
| Escherichia coli K-12 | P16528.1 | Transcriptional repressor IclR |
| Shigella flexneri | WP_011069590.1 | glyoxylate bypass operon transcriptional repressor IclR |
| Citrobacter freundii | WP_284904112.1 |  |
| Salmonella enterica subsp. salamae | ECG8596373.1 |  |
| Pseudomonas putida KT2440 | AAN68209.1 | Transcriptional regulator, IclR family |
| Azotobacter chroococcum | WP_131342702.1 | IclR family transcriptional regulator |
| Pantoea sp. Ap-967 | WP_167062202.1 |  |
| Stutzerimonas tarimensis | WP_386362031.1 |  |
| Pseudomonas putida KT2440 | AAN66998.1 | Transcription regulatory protein (pca regulon) |
| Xanthomonas sp. WHRI 1810A | WP_349801399.1 | Pca regulon transcriptional regulator PcaR |
| Azotobacter salinestris | WP_349571726.1 | IclR family transcriptional regulator |
| Stutzerimonas stutzeri | WP_213662817.1 | Pca regulon transcriptional regulator PcaR |
| Acinetobacter baylyi ADP1 | Q43992.1 | p-hydroxybenzoate hydroxylase transcriptional activator |
| Acinetobacter baumannii | WP_190595594.1 | IclR family transcriptional regulator PobR |
| Klebsiella pneumoniae | HBY6010789.1 | TPA: IclR family transcriptional regulator PobR |
| Serratia sp. S1B | PVZ88352.1 | IclR family transcriptional regulator |
| Pseudomonas putida KT2440 | AAN68217.1 | Transcriptional regulator, IclR family |
| Aeromonas caviae | GJB84029.1 | Transcriptional regulator |
| Pantoea sp. Ap-967 | WP_167062178.1 | IclR family transcriptional regulator domain-containing protein |
| Azomonas agilis | WP_144571121.1 |  |

**Supplementary Table 4. Sequence accessions used for multiple sequence alignment.** The table indicates the species name, the corresponding accession number and the associated annotation from NCBI.

**99**

**125**

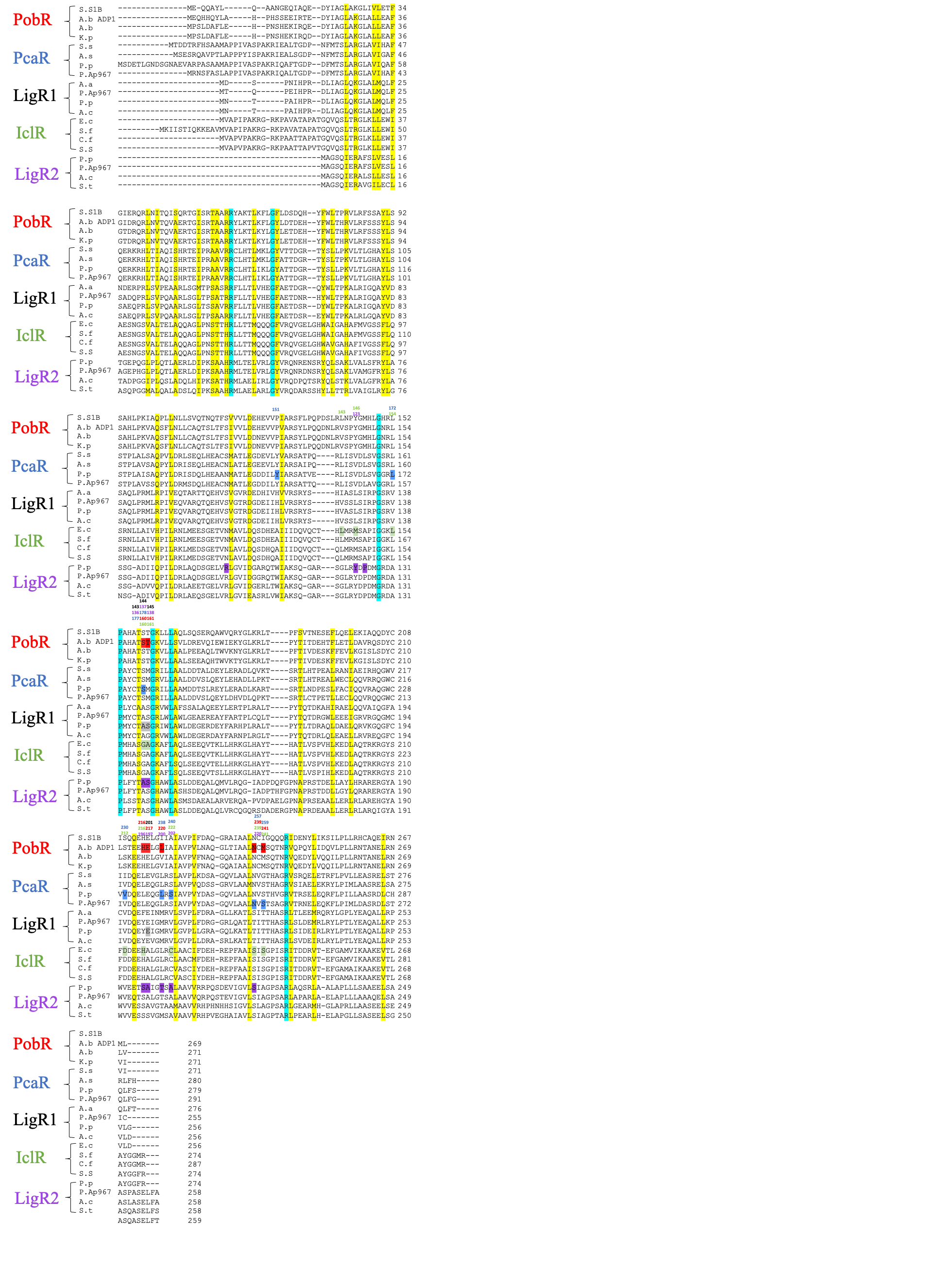

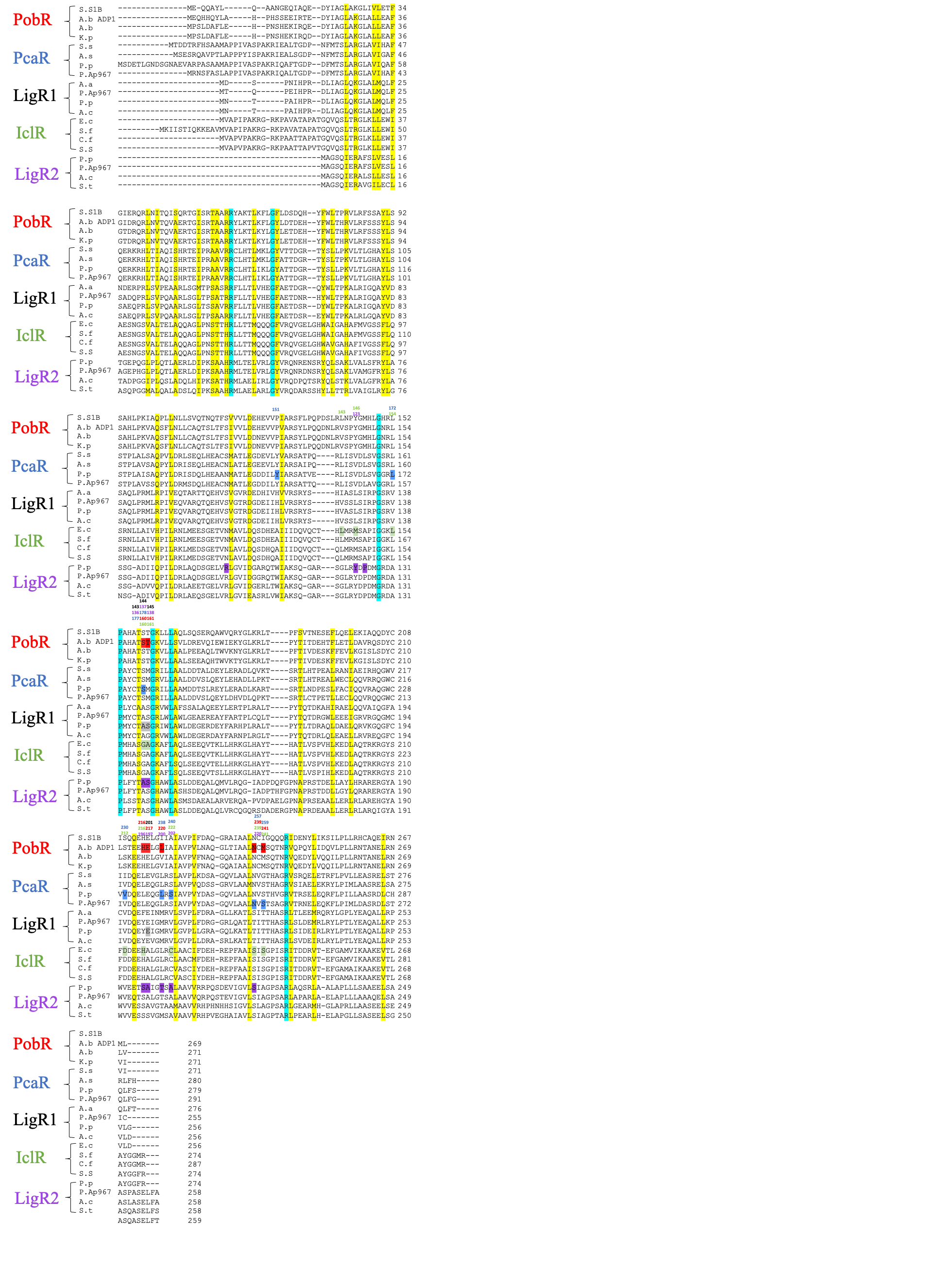
**Supplementary Figure 3. Full amino acid sequence alignments of LigR1 and LigR2 against 3 distinct families of structurally characterized IclR regulators**. Represented sequences include the full-length amino acid sequences of LigR1, LigR2, IclR, PobR, and PcaR. Sequences were aligned using the ClustalW Omega software with character counts. Columns highlighted in yellow indicates residues that exhibit weak similarity, while the teal highlighted indicate highly similar and invariant residues. The active site residues have been highlighted for each corresponding LTTR. Above each colour indicates the position of the residue in the protein.

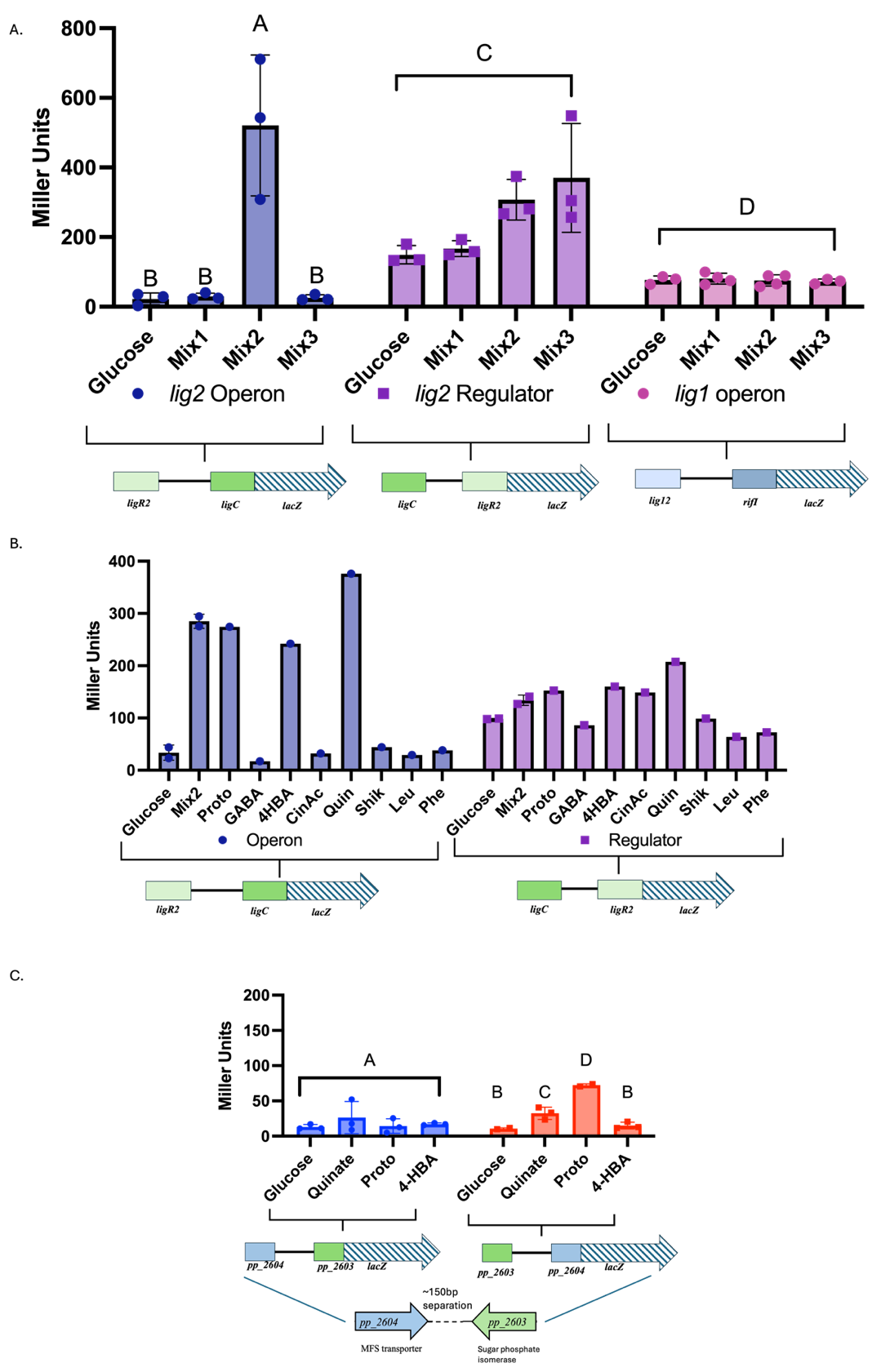

**Supplementary Figure 4. Ligand screening using 𝛽-galactosidase reporter assays.** A) 𝛽-galactosidase reporter assays of *lig2* operon expression (blue), *ligR2* expression (purple), and *lig1* operon expression (pink) with glucose, Mix1, Mix2, or Mix3. Mix1= maltose, glucose, arabinose, trehalose, fructose, xylose, mannitol, and mannose. Mix2= protocatechuate, 𝛾-butyric acid, 4-hydroxybenzoate, *t*-cinnamic acid, quinate, shikimate, leucine, and phenylalanine. Mix3= succinate, acetate, fumarate, malic acid, and oxalic acid. B) Individual Mix2 compound screen for *lig2* operon (blue) and *ligR2* (purple). Ligands were tested at a concentration of 200µM. Proto=protocatechuate, GABA= 𝛾-butyric acid, 4HBA= 4-hydroxybenzoate, CinAc= *t*-cinnamic acid, Quin= Quinate, Shik= shikimate, Leu= leucine, Phe=phenylalanine. C) 𝛽-galactosidase reporter assays of the 150bp intragenic region separating the *lig1* (red) and *lig2* (blue) operons. Replicate counts reflected by data points. Statistical analysis was conducted using a One-Way ANOVA with multiple comparisons and letters above the bars represent compact letter display of Tukey’s multiple comparisons tests. The different letters within the tested constructs represent significant differences at 95% CI. Error bars indicate the SEM.

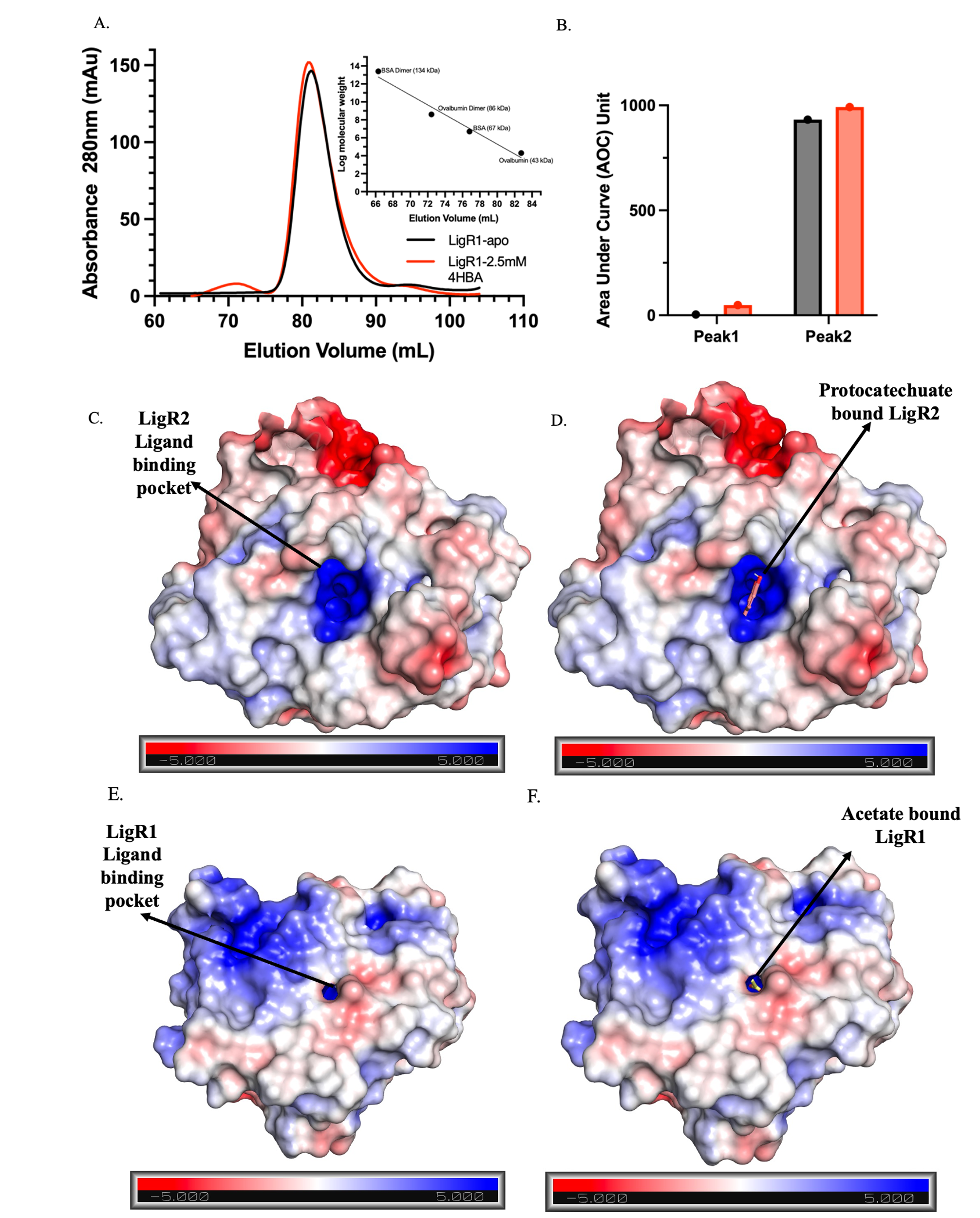

**Supplementary Figure 5. Electrostatic surface representation of LigR1 and LigR2** **A)** Gel filtration profile of LigR1 apo-protein (black) and with 2.5mM 4-hydroxybenzoate pH 7.5 (red) from 60mL to 110mL. Inserted graph shows the elution profile of bovine serum albumin (BSA) and ovalbumin standard proteins from elution volume from 65 to 85mL. Elution profiles were conducted using a HiLoad 16/60 Superdex 200pg column. **B)** Quantification of the LigR1 peaks using AUC from the elution profile C) Electrostatic surface representation of LigR2 EBD. Primary binding site is indicated with an arrow. **D)** Electrostatic surface representation of LigR2 EBD docked with protocatechuate using AutoDock. **E)** Electrostatic surface representation of LigR1 EBD. Primary binding site is indicated with an arrow. **F)** Electrostatic surface representation of the acetate bound LigR1 EBD. Positive (blue) and negative (red) electrostatic potentials are indicated by a colour gradient.

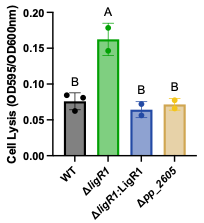

***ΔligR1 Glucose***

**Supplementary Figure 6. Cell lysis of ΔligR1 in M9 minimal media with 25mM glucose.** The supernatant of WT (black), ΔligR1(green), LigR1:ΔligR1complement (blue), and ΔligR1(brown) was collected after 26hrs of growth. The supernatant was diluted 3.33x in ¼ Bradford reagent. The OD595nm of the Bradford reagent was measured and divided by the final OD600nm of the cell to obtain the relative cell lysis. Replicate counts reflected by data points. Statistical analysis was conducted using a One-Way ANOVA with multiple comparisons and letters above the bars represent compact letter display of Tukey’s multiple comparisons tests. The different letters within the test constructs represent significant differences at 95% CI. Error bars indicate the SEM.
